# Hydrological Connectivity Enhances Fish Biodiversity in Amazonian Mining Ponds: Insights from eDNA and Traditional Sampling

**DOI:** 10.1101/2025.01.28.635072

**Authors:** Camila Timana-Mendoza, Alonso Reyes-Calderón, Patrick Venail, Ricardo Britzke, Monica C. Santa-Maria, Julio M. Araújo-Flores, Miles Silman, Luis E. Fernandez

## Abstract

The expansion of artisanal and small-scale gold mining (ASGM) in the Madre de Dios region of the Peruvian Amazon has transformed primary forests into a novel wetland complex of thousands of abandoned mining ponds. Despite their ecological relevance, post-mining recovery of these systems remains understudied, particularly regarding fish biodiversity and recolonization. In this study, we evaluate fish community richness and composition in mining ponds of different dimensions, years post abandonment, and degree of pulse flood connectivity using traditional collection-based methods and environmental DNA (eDNA) with the 12S and COI markers. We compared these two methods of biodiversity inventory and contrasted results from ASGM waterbodies with those obtained from nearby pristine oxbow lakes. Overall, we registered more fish richness at all sites using eDNA vs traditional methods, especially with the 12S marker. We identified 14 and 13 unique genera using traditional methods and eDNA, respectively, with 40 genera detected by both approaches, evidencing their complementarity. Notably, we found that the degree of pulse flooding connectivity was the main predictor of species richness among the abandoned mining ponds (p-value < 0.05). We registered 11 to 22, 23 to 71, and 56 morphospecies in non-flooded mining ponds, pulse flooded mining ponds and nearby oxbow lakes, respectively. Furthermore, the fish community composition of mining ponds most influenced by pulse flooding were similar to that of pristine lakes. Our findings highlight the role of hydrological connectivity in ecological recovery within mining-impacted wetlands. Future restoration efforts should enhance aquatic connectivity to accelerate recovery in post-mining environments.

## Introduction

The Amazon basin is recognized for its exceptional biological diversity and indigenous communities, standing as one of the most vital and ecologically significant regions on Earth (Cassemiro et al., 2023; Heilpern et al., 2022). Notwithstanding its ecological relevance, artisanal and small-scale gold mining (ASGM) is occurring in many regions within the Amazon, severely affecting the environment as well as the economy, public health, and contributing to local and transnational crime (Dethier et al., 2023a; Esdaile and Chalker, 2018; Gasparinnetti et al., 2024; Keane et al., 2023). ASGM, which typically occurs along rivers and primary forests, is a significant driver of the extensive and ongoing deforestation within the Amazon basin, increases in sediment loads in rivers, water siltation, mercury pollution (used for gold amalgamation), and the alteration of hydro-geological processes, among other negative effects (Alvarez-Berrios & Aide, 2015; Cheng et al., 2023; Crespo-Lopez et al., 2021; Dethier et al., 2023a; Swenson et al., 2011; Veiga et al., 2006).

Within the Amazon basin, the Madre de Dios Region in Perú is a biodiversity hotspot that has been particularly impacted by ASGM activities (Asner & Tupayachi, 2016; Barocas et al., 2023; Caballero et al., 2018; Dethier et al., 2019; Diringer et al., 2020; Gerson et al., 2020; Kahhat et al., 2019; Martinez et al., 2018; Paiva et al., 2023; Swenson et al., 2011). Mining activities in Madre de Dios have resulted in more than 120,000 hectares of primary forest cleared and transformed into an artificial wetland system and bare or naturally revegetating land (Caballero et al., 2018; Caballero et al., 2020; Camalan et al., 2022; Dethier et al., 2023a) where, due to extensive soil degradation and uncertain land tenure, reforestation efforts have been difficult to deploy (Román-Dañobeytia et al., 2021; Garate-Quispe et al., 2021; Herrera-Machaca et al., 2024). The Madre de Dios post-mining landscape presents thousands of small mining pits or ponds ranging from 0.1 to 28.8 hectares in size, created to extract alluvial deposits of gold (Alvarez-Berrios & Aide, 2015; Asner & Tupayachi, 2016; Caballero et al., 2020). These ponds are frequently left abandoned, and represent a significant risk due to their potential for mercury methylation (Gerson et al., 2020; Gerson et al., 2021). Aquatic communities have been able to recolonize these new artificial ecosystems, facilitated by their connectivity with rivers and streams during the rainy season (Araujo-Flores et al., 2021). Annual river flood pulses during the rainy season serve as a main driver of species accumulation in a river floodplain (Baumgartner et al., 2018; Melack & Coe, 2021; Pereira et al., 2017), by providing them with nutrients and individuals (Araujo-Flores et al., 2021; Fernandes et al., 2014; Strahler, 1952). In addition, isolated mining ponds may support aquatic communities through fish introductions by birds, human stocking, or occasional climate-driven connections (Cooke et al., 2022; Green et al., 2023). In any case, post-mining recovery and aquatic communities’ recolonization of these systems remain understudied, particularly concerning fish biodiversity.

To improve restoration and management strategies in these aquatic environments, extensive and continuous monitoring is necessary, to assess their dynamics and behavior over time and space (Deiner et al., 2017; Kelly et al., 2014; Magurran et al., 2010). Traditional methods, based on species collection through nets and taxonomic identification using morphological keys present limitations such as cost, time, labor, and the need of taxonomic reference collections that are sparse or nonexistent (Blattner et al., 2021; Cristescu & Hebert, 2018; Evans et al., 2017; van der Sleen & Albert, 2017; Wilcox et al., 2013). Alternatively, environmental DNA (eDNA), which refers to the detection of genetic material left by organisms in the sampled environment (Deiner et al., 2017) has proven to be a complementary method in wildlife research in general (Keck et al., 2022; Klymus et al., 2017; Munian et al., 2024; Ruppert et al., 2019; Takahashi et al., 2023; Veron et al., 2023), and in fish communities in the Neotropics in particular (Cilleros et al., 2019; Coutant et al., 2023; de Santana et al., 2021; Jackman et al., 2021; Mariac et al., 2022; Sales et al., 2021), including in this post-mining environment (Timana-Mendoza et al., 2024). Particularly, eDNA has the potential to be a powerful tool for gathering aquatic biodiversity data across extensive aquatic landscapes that are difficult to access and survey (Carvalho et al., 2021; Harper et al., 2019; Takahashi et al., 2023), though its ability to capture total aquatic biodiversity, or even total fish biodiversity, remains to be tested, as does the simple congruence between eDNA-based and collections-based surveys.

In this study, we assessed the richness and composition of fish communities in mining ponds of varied surface areas and years post abandonment, affected by various degrees of seasonal flooding, within the Madre de Dios Region in Southern Peru. Our aim was to depict what are the main factors impacting the recolonization dynamics in these novel ecosystems. We used both eDNA and traditional biological monitoring methods simultaneously, and looked into the taxonomic resolution of two DNA markers commonly used for eDNA analysis of ichthyofauna. We included nearby unmined oxbow lakes in the study, to serve as local reference for richness and community composition.

## Materials and methods

### Study area

We evaluated nine abandoned mining ponds and two nearby unmined oxbow lakes within the Madre de Dios River basin, located in the Madre de Dios region, at the headwaters of the tropical Amazon and the foothills of the Andes (11°37′46.60″S, 70°32′25.84″O) in southeastern Peru (Figure 1). In this area, annual precipitation varies from 1700 to 3300 mm per year, depending on proximity to the Andes (Torre-Marin et al., 2021). Intense local precipitation and poor drainage result in seasonally recurrent short-duration fluvial floods (often <1–2 weeks), generating floodplains between December and April. These changing conditions contribute to the Madre de Dios River basin unique ecological importance related to its high fish diversity (ca. 700 sp.), and for fish spawning migrations within the Amazon system (Araújo-Flores et al., 2021).

**Figure 1.**
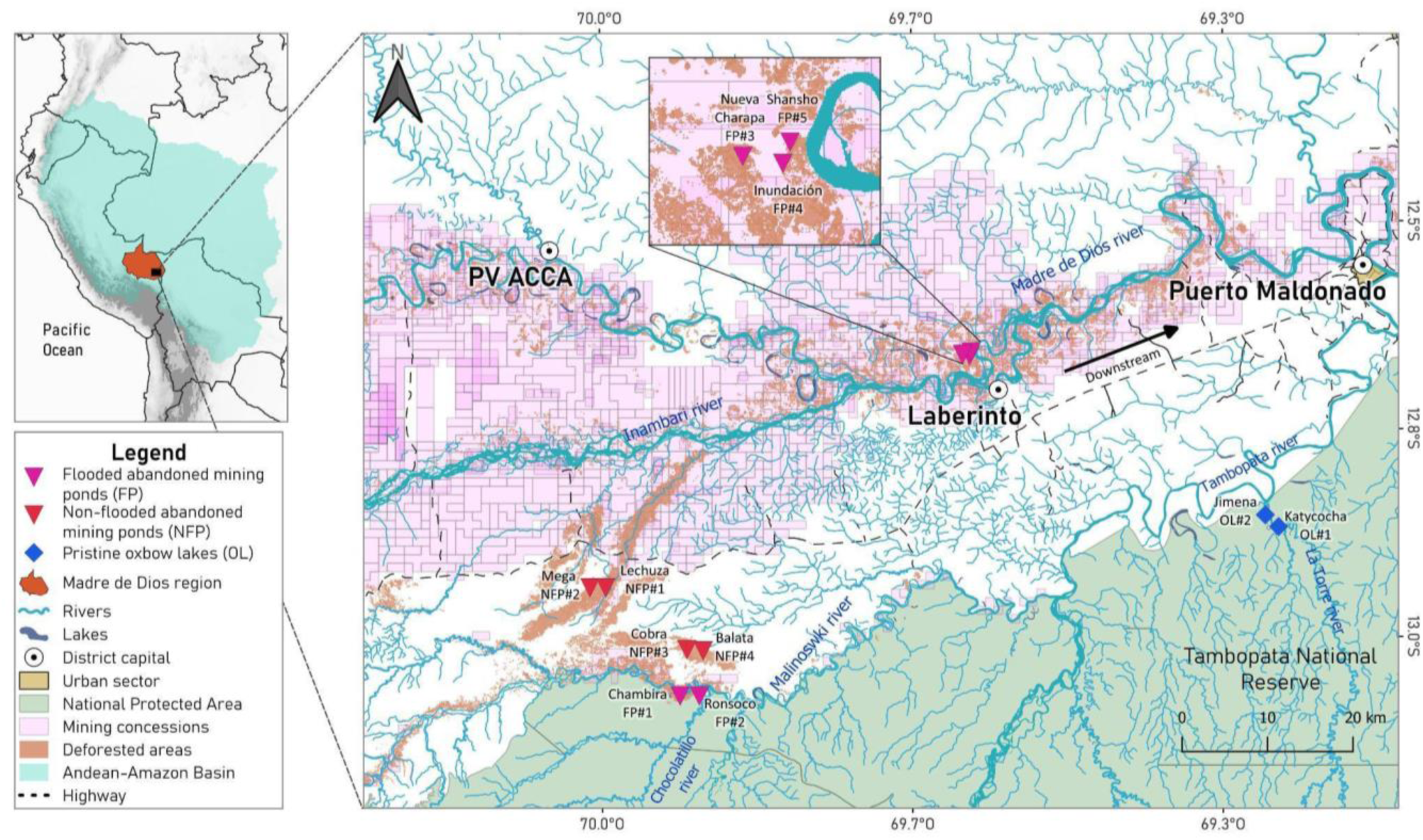
Study area within the Madre de Dios River basin showing the 11 lentic systems surveyed in the study: five flooded abandoned mining ponds (FP), four non-flooded abandoned mining ponds (NFP) and two pristine oxbow lakes (OL).

Among the mining ponds surveyed, five belong to a floodplain ecosystem, and four to a recently deforested *terra firme* or non-flooded ecosystem (Figure 1). The mining ponds within the floodplain ecosystem (flooded [abandoned mining] ponds, FP) are influenced by the nearby rivers ‘Madre de Dios’ and ‘Malinowski’ through rainy season flooding (see Figure 1 and Table 1). In contrast, the abandoned mining ponds within the *terra firme* (non-flooded abandoned mining ponds, NFP) are located inside the mining area known as “La Pampa”, south of the Interoceanic Highway (12°59′26.88′′S, 69°56′36.42′′O) and are either not influenced by any streams or slightly influenced by small and oligotrophic Amazonian blackwater streams tightly linked with local rainfall regimes (see Figure 1 and Table 1). Flood zones were delimited using ArcGIS Pro 2.8.7 and a Digital Elevation Model (DEM). Elevation values (‘Z’) and geomorphological criteria were analyzed to identify flood levels in the two floodplain ecosystem study sites: 243-248 m.a.s.l. in the FP#1 and FP#2 within the ‘PVC Azul’ area, and 215-220 m.a.s.l. in the FP#3, FP#4 and FP#5 within the ‘Laberinto’ area (Figure 1). These results were validated with high-resolution PlanetScope images (3m) captured during the seasonal flooding peak, in February. Additionally, the abandonment date was estimated through a multitemporal analysis of Landsat 8 satellite images using supervised classification algorithms to determine the last documented mining impact in each area. This approach enabled tracking land cover disturbances and establishing abandonment periods up to the evaluation date. The two nearby unmined oxbow lakes (OL) - used as reference - are located within the Tambopata National Reserve (see Figure 1 and Table 1). All these lentic systems were sampled during the dry season transition from May to June 2022 to avoid recent contributions from lotic systems.

**Table 1.**
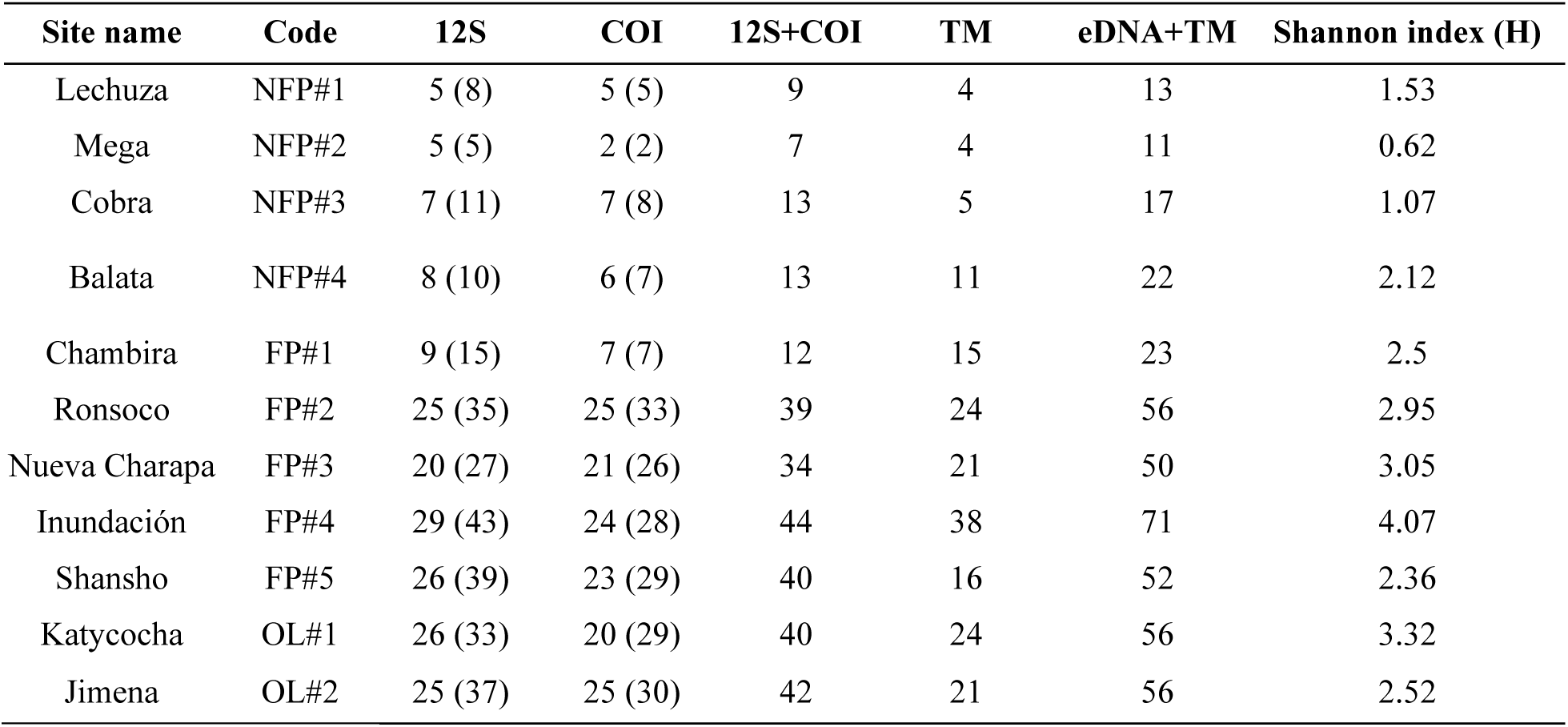
Main characteristics of the abandoned mining ponds and unmined oxbow lakes surveyed in the study. NFP: Non-flooded [abandoned mining] ponds. FP: Flooded [abandoned mining] ponds. OL: [pristine] Oxbow lakes.

### eDNA sampling and purification

We collected water samples for eDNA analysis at each site as follows: We took three to five 4L water samples across the mining pond or oxbow lake using a clean 4L plastic bucket and a small boat. These 4L water samples were pooled into a clean 20L plastic bucket to make a single composite sample. Two such composite samples were generated for each site surveyed. Both the 4L and the 20L buckets used to collect environmental samples were pre-cleaned by washing them with commercial powder detergent, rinsed thoroughly with tap water, then treated by submerging them in 1% v/v chlorine bleach solution for at least 1 hour, and then rinsed thoroughly with bottled drinking water. Buckets were rinsed with the pond or lake’s water at least three times before collecting the water samples used for the analysis. From each 20L composite sample, we took a 1L water subsample with a clean 1L high-density polyethylene (HDPE) plastic bottle. These bottles were previously cleaned by washing them with commercial detergent, thoroughly rinsed with tap water, submerged in 1% v/v chlorine bleach solution for at least 1 hour, rinsed thoroughly with distilled water (Direct-Q 3 UV, Merck Millipore, Massachusetts, US), let dry in a UV-irradiated bench, and finally irradiated 30 min with UV-C light. They were packed in a clean cabinet to be opened only in the field for sample collection. We obtained two 1L subsamples per site (one from each 20L composite sample) to make a total of 22 environmental water samples from the 11 sites surveyed. The 1L plastic bottles were then stored in a cooler at <8°C and transported to the field station, where they were immediately filtered (within 6 to 12 h after sample collection). We filtered 400 mL to 950 mL from this 1L subsamples (see supplementary information 1) through a glass fiber membrane of 1.2 μm pore size (Merck Millipore, Massachusetts, US) using a MF5Pro Magnetic Filter Holder and Funnel (Rocker Scientific Co., Taiwan) with a vacuum pump (KNF Group, Freiburg, Germany), until the filter was clogged. We then passed 5 mL pure ethanol (Sigma-Aldrich, Missouri, US) through the filters and stored moist filters in 5 mL centrifuge tubes (Eppendorf, Hamburg, Germany). A field station control was obtained by filtering bottled drinking water alongside the environmental water samples, to assay for possible environmental DNA contamination in the field station.

We obtained total environmental DNA using the NucleoSpin eDNA Water Kit (Macherey-Nagel, Düren, Germany), following manufacturer guidelines without modifications. We eluted total DNA in 100 μL Elution Buffer, provided in the kit. We performed eDNA extractions in a DNA extraction chamber dedicated to eDNA work. Before each extraction batch, we carefully wiped the bench surface, pipettes, scissors, tweezers, and all other equipment using DNA/RNA-ExitusPlus™ (PanReac AppliChem) followed by ultraviolet exposure for 30 min. We used filtered tips for all steps involving pipetting. We included DNA extraction blanks, consisting of ultra-pure water (Direct-Q 3 UV, Merck Millipore) processed alongside the samples, as laboratory negative controls. Total DNA samples were stored at −20 °C until further processing.

### Library preparation and sequencing

We prepared eDNA libraries targeting regions within the mitochondrial cytochrome oxidase I gene (COI) and the mitochondrial 12S ribosomal RNA (12S rRNA) gene, using the Illumina MiSeq dual-barcoded two-step PCR amplicon sequencing protocol, following guidelines described in the Illumina 16S Metagenomic Sequencing Library Preparation Manual (Illumina Technology, 2013), with the following modifications: The COI region was PCR-amplified using the primer set MK1-F1 5’-TCGTCGGCAGCGTCAGATGTGTATAAGAGACAG-TCHACHAAYCAYAAAGAYATYGGYACYCT-3’ and MK1-R1 5’-GTCTCGTGGGCTCGGAGATGTGTATAAGAGACAG-ACYATRAARAARATYATYACRAADGC described in Mariac et al. (2022), which yields 185 bp fragments (including primers). Four PCR reaction replicates were set per eDNA sample in 11 µL total reaction volume. PCR reactions contained 1X DreamTaq Green PCR Master Mix (ThermoFisher Scientific, Waltham, Massachusetts, USA), 0.4 μM of each primer, 0.8 mg/mL BSA, 3 mM MgCl_2_ and 1 µL template DNA. Amplification parameters included an initial denaturation at 95°C for 3 min, followed by 40 cycles of 95°C for 30 s, 51°C for 30 s and 72°C for 1 min, and a final extension step of 72°C for 5 min. The 12S region was PCR-amplified using the primer set F1 5’-TCGTCGGCAGCGTCAGATGTGTATAAGAGACAG-ACTGGGATTAGATACCCC-3’ and R1 5’-GTCTCGTGGGCTCGGAGATGTGTATAAGAGACAG-TAGAACAGGCTCCTCTAG-3’ described in Kelly et al. (2014) (Riaz et al., 2011 in Kelly et al., 2014). 12 PCR reaction replicates were set for each eDNA sample, in 8 µL total reaction volume. PCR reactions contained 1X DreamTaq Green PCR Master Mix, 0.4 μM of each primer, 0.8 mg/mL BSA, 1.5 mM MgCl_2_, 3% DMSO and 0.9 μL template DNA. Amplification parameters included an initial denaturation at 95°C for 15s, followed by 45 cycles of 95°C for 15 s, 57°C for 30 s, and 72°C for 30 s, and a final extension step of 72°C for 5 min (Timana-Mendoza et al., 2024). In both cases, PCR reaction replicates per eDNA sample were pooled and verified by agarose gel electrophoresis using 5 µL of pooled PCR product. Pooled amplification products were purified using sample purification beads (SPB, Illumina, San Diego, California, USA). Index PCR reactions were carried out using 1x DreamTaq Green PCR Master Mix, 2 μL of clean amplicon PCR product as a template, and indexes from the Nextera XT Index Kit v2 (Illumina) in a final volume of 20 μL. Index PCR amplification parameters included an initial denaturation at 95°C for 3 min followed by 8 cycles of 95°C for 30 s, 55°C for 30 s, and 72°C for 30 s, and a final extension step of 72°C for 5 min. Index-PCR products were analyzed by agarose gel electrophoresis and cleaned using the same SPB procedure described above. Samples were then quantified using the Qubit fluorometer 4 (Invitrogen, Waltham, Massachusetts, USA) with the Qubit dsDNA HS kit (Invitrogen), and then normalized by diluting all indexed libraries to 4 nM in nuclease-free water. Libraries were denatured and sequenced at 8 pM final concentration (20% Phi-X control added) in the Illumina MiSeq sequencing system using the MiSeq Reagent Kit v3 (2×300 bp) following the standard run protocol (Illumina).

### Bioinformatic analysis

Bioinformatic analysis was performed via a custom pipeline from NatureMetrics, UK. In brief, sequences were demultiplexed with bcl2fastq based on the combination of the i5 and i7 index tags. Paired-end FASTQ reads for each sample were merged with USEARCH (Edgar, 2010) requiring a minimum of 80% agreement in the overlap. Forward and reverse primers were trimmed from the merged sequences using cutadapt (Martin, 2011) with a length filter of 125-135bp for COI primer and 80-120bp for 12S primer (post primer removal). Sequences were quality filtered with USEARCH to retain only those with an expected error rate per base of 0.01 or below and dereplicated by sample, retaining singletons. Unique sequences from all samples were denoised in a single analysis with UNOISE (Edgar, 2016) requiring retained zOTUs (zero-radius Operational Taxonomic Units) to have a minimum abundance of 8 in at least one sample. Consensus taxonomic assignments were made for each zOTU using sequence similarity searches against NCBI nucleotide (NCBI nt) for 12S primer, and NCBI and BOLD (Ratnasingham and Hebert, 2007) for COI primer. Searches against databases were made using blastn (Altschul et al., 1990; Camacho et al., 2009) and required hits to have a minimum e-score of 1e-20 and cover at least 90% of the query sequence. The taxonomic identification associated with all hits was converted to match the Global Biodiversity Information Facility (GBIF) taxonomic backbone. Assignments were made to the lowest possible taxonomic level where there was consistency in the matches, with minimum similarity thresholds of 99%, 97%, and 95% for species, genus, and higher-level assignments, respectively. zOTUs were clustered at 97% similarity with USEARCH to obtain OTUs. Automated identifications were sense-checked against GBIF occurrence records for presence in the sampling country, and elevated to higher taxonomic levels where required (rgbif; Chamberlain et al., 2023), as well as the CAS Eschmeyer’s Catalog of Fishes database (Eschmeyer et al., 2024), and the reports in Peru (Chuctaya et al., 2022; Meza-Vargas et al., 2021; Ortega et al., 2012).

Following taxonomic assignment, zOTUs were clustered into OTUs to minimize the number of sequence variants for a species (that may be present due to intraspecific variations, or amplification or sequencing artifacts). Supervised clustering was done using a combination of USEARCH UPARSE (Edgar, 2013) and a custom pipeline that takes into account sequence similarity, co-occurrence patterns, abundance profiles, and taxonomy to prevent the over-clustering of distinct, closely related species. Chimeric sequences were excluded, and an OTU-by-sample table was generated by mapping all dereplicated reads for each sample to the OTU representative sequences with USEARCH at an identity threshold of 97%. The OTU table was filtered to remove low abundance OTUs from each sample. The minimum read count for a detection of an OTU within a sample at approximately 20 reads. To do this the percentage threshold across all samples was identified within the dataset that most closely achieved this and applied this threshold across all samples. Unassigned OTUs, and OTUs identified to human and domesticated mammals, were removed from the dataset for subsequent analyses.

### Traditional fish sampling

Fish samples were collected at the same time as eDNA samples or within two weeks after eDNA sample collection. We used a 10 mm seine net (30 m long and 2 m high with a 0.6 cm mesh opening) and a gill net of 0.4 mm nylon (40 m long and 2 m high composed of three nets of mesh sizes 5, 7.5 and 10 mm) to catch individual fish. Sampling with the seine net was done three times at three sites (randomly picked) within each lentic system, starting from the middle of the waterbody towards the shoreline (Araújo-Flores et al., 2021). The gill net was employed overnight (sunset to sunrise) and placed parallel to the shore. Individual fish were identified at the species level when possible and otherwise to the genus level using morphological observations and a variety of keys and checklists together with the nomenclatural assignments of Eschmeyer et al. (2024).

### Data analysis

To analyze species richness among the sites surveyed we compared taxonomic groups at the morphospecies level, that is, unique MOTUS mapped at the genus or species level, when available, for eDNA, and unique morphological assignments when using traditional methods. The data set used in the analysis, listing the number of reads mapped to a given order, family, genera and species per sampling site is presented as supplementary information 2 for the 12S marker, and as supplementary information 3 for the COI marker.

When using the COI marker, there were four instances where more than one species was mapped to a unique MOTU with 100% similarity. In these cases, the species reported for Peru (Meza-Vargas et al., 2021; Chuctaya et al., 2022; Ortega et al., 2012; Eschmeyer et al., 2024) among the mapped set was selected. This was done as follows: Between *Astyanax bimaculatus* and *Psellogrammus kennedyi*, *Astyanax bimaculatus* was selected. Between *Prochilodus nigricans* and *Prochilodus rubrotaeniatus*, *Prochilodus nigricans* was selected. *Between Hypoptopoma gulare* and *Hypoptopoma steindachneri, Hypoptopoma gulare* was selected. However, between *Auchenipterus ambyiacus* and *Auchenipterus nuchalis*, both species have been reported for Peru, so this unique MOTU was taken as a single morphospecies referred to as *Auchenipterus ambyiacus / Auchenipterus nuchalis* in the analysis. For other MOTUs’ assignments to fish not reported for Peru, we modified the taxonomic designations as follows: *Acestrorhynchus lacustris* (99.06% similarity) was relabeled as A. aff. *lacustris*, *Leporinus friderici* (100% similarity) as L. aff. *friderici*, and *Eigenmannia trilineata* (99.21% similarity) as E. gr. *trilineata* (see Table 3).

To assess the effect of different ecological contexts or lentic system characteristics (flooding status, year of abandonment and pond dimensions) on species richness among the sites surveyed, we performed a multifactorial Analysis of Variance (ANOVA). The analysis was performed using the *‘aov’* function from the *‘stats’* package (R Core Team, 2024). Then, to assess the effect of flooding on the composition of fish communities between the three conditions described previously: flooded ponds (FP), not flooded ponds (NFP) and pristine oxbow lakes (OL), we used the Jaccard dissimilarity index (Baselga, 2012). Briefly, we created a presence-absence matrix comprising all identified morphospecies by traditional methods and eDNA (considering both the 12S and COI markers) (Table 3). We, then, ran the Jaccard dissimilarity index between sites using the ‘*vegdist*’ function from the ‘*vegan*’ package (Oksanen et al., 2023), and performed a Principal Coordinate Analysis (PCoA) to visualize the patterns in two-dimensional plots using the Jaccard dissimilarity matrix with the *’cmdscale’* function from the *‘stats’* package (R Core Team, 2024), extracting two-dimensional ordination scores. We then tested for population assemblages’ similarities between the three conditions (FP, NFP, OL) using a permutational multivariate analysis of variance (PERMANOVA), comparing assemblage variation with an analysis of homogeneity of multivariate dispersion using the *‘adonis’* function from the *‘vegan’* package (Oksanen et al., 2023). The significance was assessed with 999 permutations. All data analyses were conducted in R version 4.3.1., using RStudio 2023.06.2 as the front end (Allaire, 2012), and Python 3.9 (Van Rossum, 2019).

## Results

### Contrast between the 12S and COI markers for eDNA analysis

Contrasting results were obtained when using either the 12S or the COI marker for eDNA analysis. With the 12S marker, we obtained 2,588,108 processed reads, of which 1,617,483 were mapped to class Actinopterygii, with an average of 147,043 reads per sample (range = 1,012 - 235,431). With the COI marker we obtained 1,783,519 processed reads, of which only 218,614 were mapped to class Actinopterygii, with an average of 19,874 reads per sample (range = 1,160 - 80,732). In relation to taxa identification, with the 12S marker we obtained 63 MOTUs; all of which were mapped at the order level, 61 at the family level, 43 at the genera level, and only 20 at the species level. They belong to 5 orders, 21 families, 39 genera, and were grouped into 41 morphospecies (i.e., MOTUs mapped at the genus or species level, when available) (see supplementary information 2). In contrast, with the COI marker we obtained 61 MOTUs; all of which were mapped at the order level, 58 at the family level, 46 at the genera level, and 30 at the species level. They belong to 4 orders, 16 families, 38 genera, and were grouped into 44 morphospecies (see supplementary information 3). Overall, the 12S marker registered more MOTUs than the COI marker at all sites (Figure 2). At the morphospecies level, comparable results were obtained across sites using either eDNA marker and traditional monitoring (Table 2). Pertaining to the eDNA markers used, at the family level, the 12S marker identified five families not recorded by the COI marker, whereas the COI marker identified only one family not recorded by the 12S marker. At the genera level, the 12S marker identified 15 genera not recorded by the COI marker, whereas the COI marker recorded 14 genera not recorded by the 12S marker. At the morphospecies level, both markers shared 15 morphospecies, the 12S marker identified 26 non-shared morphospecies and the COI marker identified 29 non-shared morphospecies; while 20 and 15 MOTUs could not be identified at the morphospecies level with the 12S and COI markers, respectively (Figure 3). At the species level, the COI marker allowed us to identify 10 more species than the 12S marker considering all sites surveyed (supplementary information 3).

**Figure 2.**
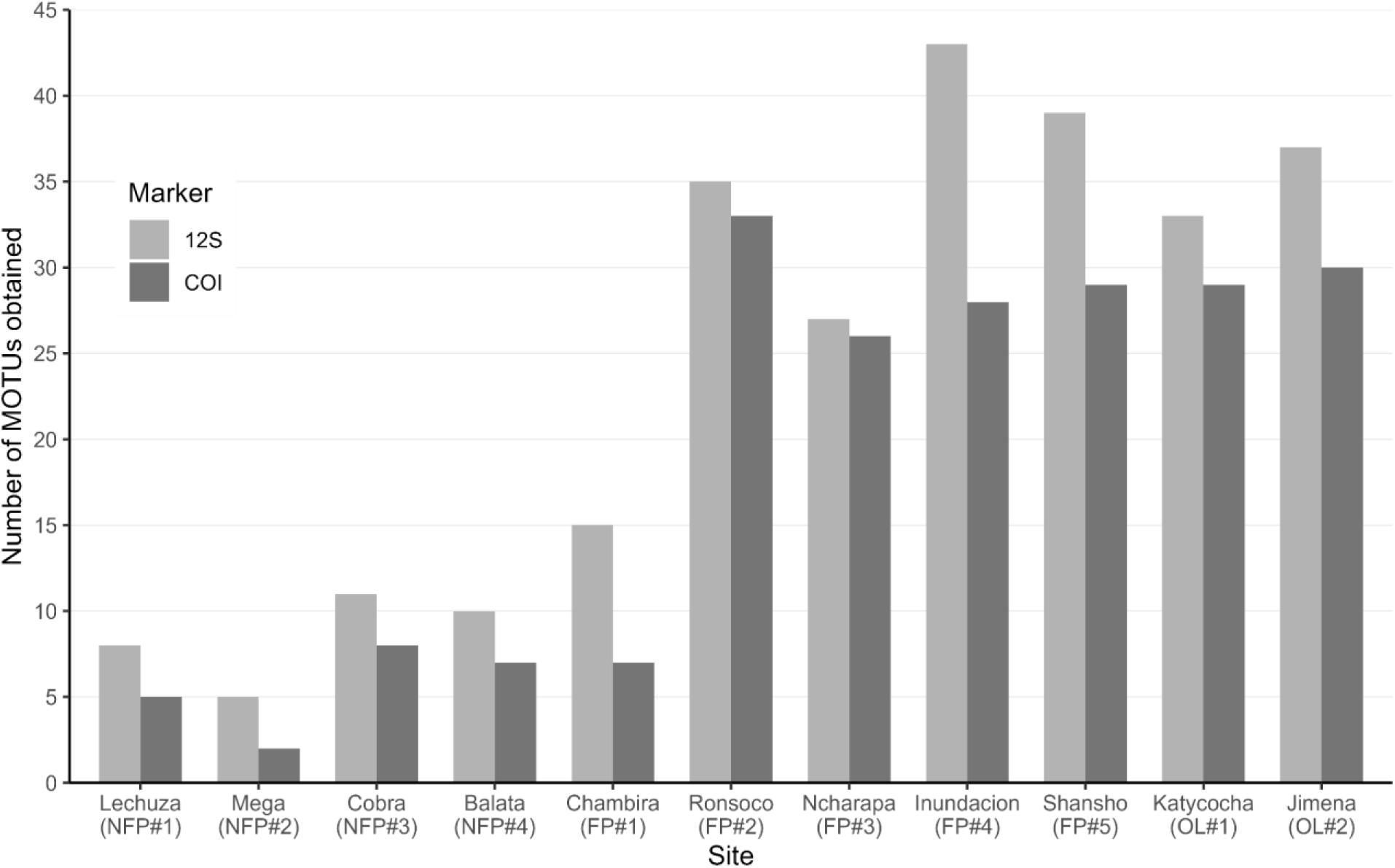
Number of MOTUs belonging to Actinopterygii obtained using the 12S (Riaz et al., 2011 in Kelly et al., 2014) and COI (Mariac et al., 2022) markers in the abandoned mining ponds and nearby pristine oxbow lakes surveyed in the study. NFP: Non-flooded [abandoned mining] ponds. FP: Flooded [abandoned mining] ponds. OL: [pristine] Oxbow lakes.

**Figure 3.**
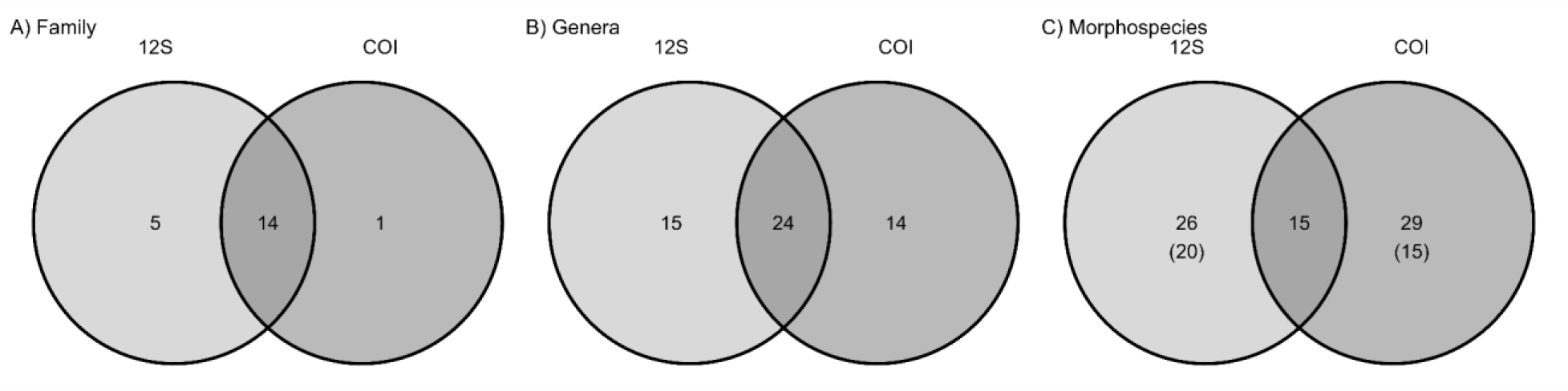
Venn diagrams showing the shared (and not shared) number of families (A), genera (B) and morphospecies (C) by the 12S (Riaz et al., 2011 in Kelly et al., 2014) and COI (Mariac et al., 2022) markers across all sites. Numbers in parenthesis are MOTUS not mapped at morphospecies level.

**Table 2.**
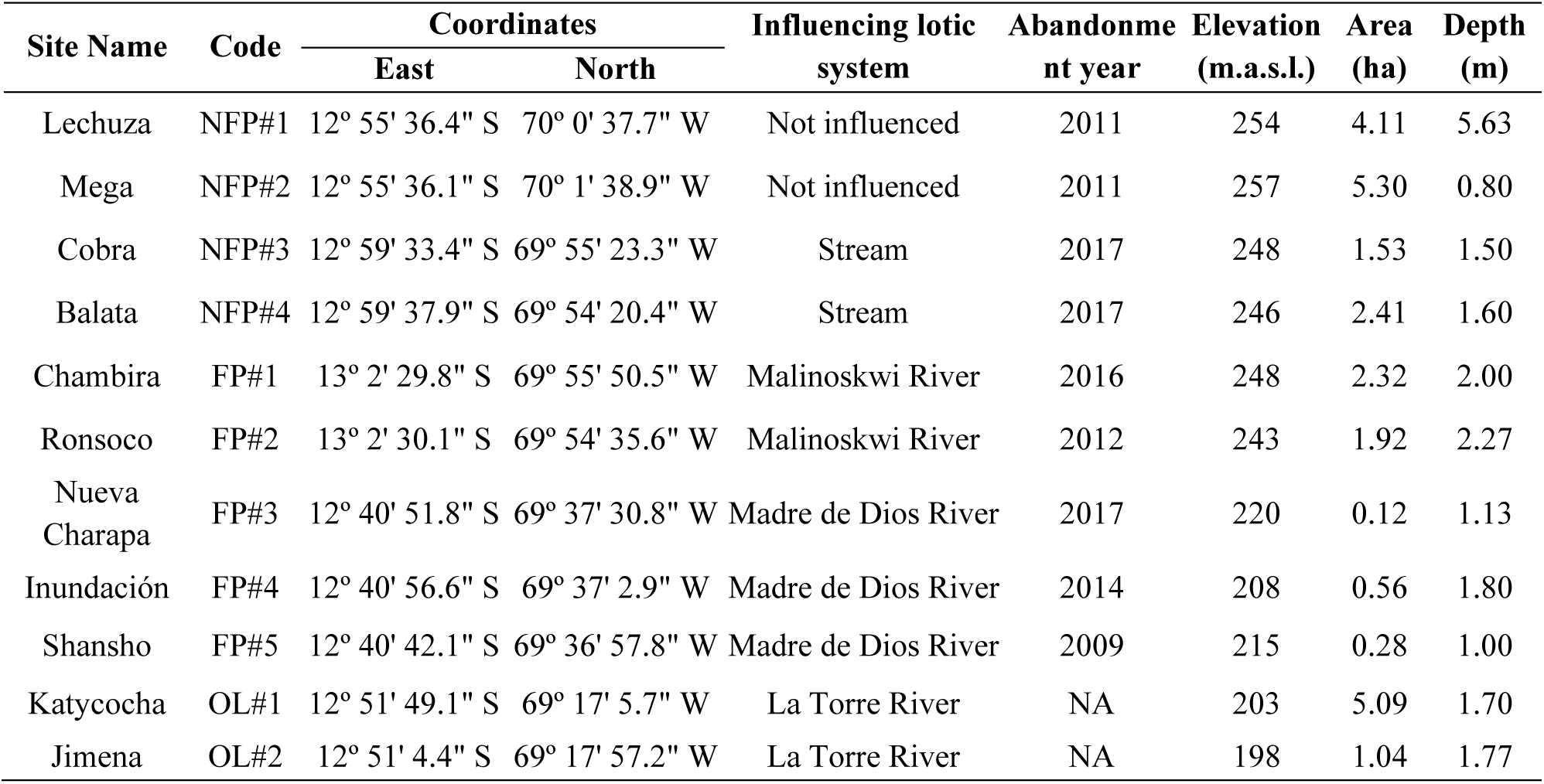
Number of morphospecies identified with the 12S marker, the COI marker, both markers combined (12S+COI), traditional methods (TM) and both eDNA and traditional methods combined (eDNA+TM). Numbers in parentheses represent the total number of MOTUs for each eDNA marker. The Shannon-Wiener diversity index (H) shown was calculated using the data obtained with the traditional sampling method only. NFP: Non-flooded [abandoned mining] ponds. FP: Flooded [abandoned mining] ponds. OL: [pristine] Oxbow lakes.

Differences in fish community composition were observed across the sites surveyed when using either DNA marker (Figure 4). The 12S marker identified more families than the COI marker at all sites (supplementary information 4), and several families were exclusively detected by the 12S marker (Figure 3): Acestrorhynchidae, Iguanodectidae (found only in Balata NFP#4 and Cobra NFP#3), Parodontidae, Synbranchidae (found only in Jimena OL#2), and Heptapteridae (found only in Inundación FP#4). In contrast, the COI marker was able to identify the Triportheidae family, not registered when using the 12S marker. In addition, the 12S marker was unable to resolve (i.e., assign to lower taxonomic levels) several MOTUs mapped at the order Characiformes in Lechuza NFP#1, Cobra NFP#3, Balata NFP#4 and Chambira FP#1, where the COI marker did not show any unassigned MOTUs at the family level. Both markers consistently detected the family Cichlidae across all sites (Figure 4, supplementary information 4). Similarly, the families Erythrinidae and Characidae were present at all locations except for Mega NFP#2. At this site neither molecular marker registered these families, although traditional sampling did confirm the presence of *Hoplias malabaricus* from the family Erythrinidae.

**Figure 4.**
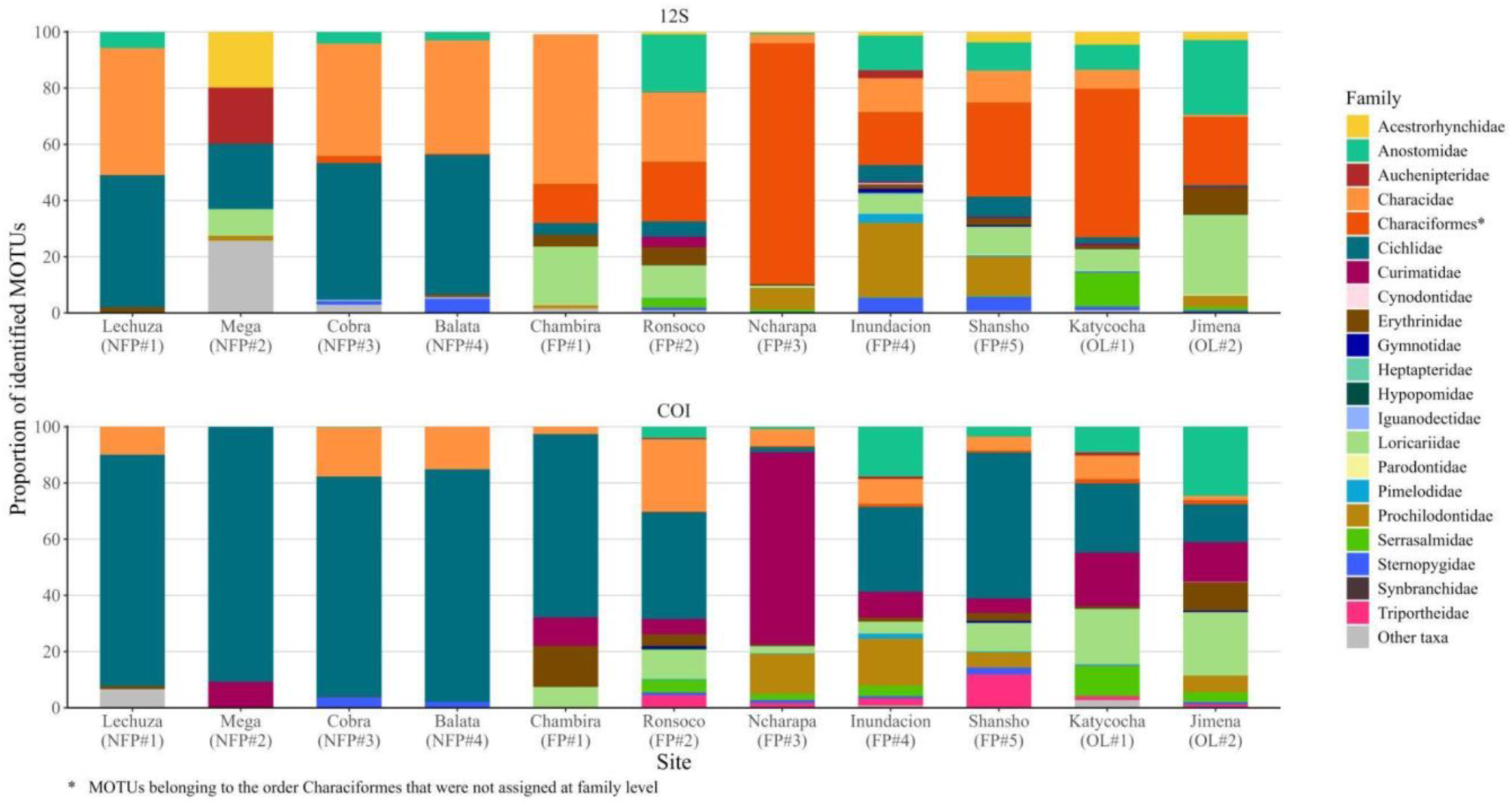
Family composition of fish communities in the abandoned mining ponds and oxbow lakes identified with eDNA, using the 12S marker (Riaz et al., 2011 in Kelly et al., 2014, top panel) and the COI marker (Mariac et al., 2022, bottom panel). Other taxa, represented by gray color (birds, amphibians, and reptiles), were identified at all sites, although at lower proportions (see supplementary information 6 and 7). NFP: Non-flooded [abandoned mining] ponds. FP: Flooded [abandoned mining] ponds. OL: [pristine] Oxbow lakes.

Overall, we observed lower family diversity at the non-flooded mining ponds (NFP) compared to flooded ponds (FP) and oxbow lakes (OL), with a predominance of the families Cichlidae, Characidae, and Curimatidae (Figure 4, supplementary information 4).

### Contrast between eDNA and traditional methods for biodiversity monitoring

Considering all the sites surveyed, we detected a total of 123 morphospecies using eDNA and traditional methods combined, which comprised 68 genera, 26 families, and 6 orders (Table 3). With the eDNA approach, using both markers combined, we identified a total of 70 morphospecies, belonging to 54 genera, 21 families and 5 orders. In contrast, when using traditional methods, we recorded 76 morphospecies belonging to 53 genera, 20 families and 6 orders (see supplementary information 5). Notably, at the species level, the traditional methods identified 61 species, whereas the eDNA method identified 41 species with 43 and 31 unassigned MOTUs at the species level with the 12S and COI marker, respectively (Table 3).

**Table 3.** Prevalence of morphospecies registered by eDNA (12S and COI) and traditional methods (TM) among the 11 sites surveyed. Values in the table indicate the number of sites where a given morphospecies was encountered using either method.

| Family | Genus | Species | eDNA |  | TM |
| --- | --- | --- | --- | --- | --- |
|  |  |  | 12S | COI |  |
| Acestrorhynchidae | *Acestrorhynchus* | *Acestrorhynchus* aff. *lacustris* | 7 | 0 | 0 |
| Acestrorhynchidae | *Acestrorhynchus* | *Acestrorhynchus falcatus* | 0 | 0 | 4 |
| Anostomidae | *Leporinus* | *Leporinus* aff. *friderici* | 0 | 5 | 0 |
| Anostomidae | *Leporinus* | *Leporinus* cf. *subniger* | 0 | 0 | 6 |
| Anostomidae | *Leporinus* | *Leporinus* sp. | 9 | 2 | 0 |
| Anostomidae | *Megaleporinus* | *Megaleporinus trifasciatus* | 0 | 3 | 0 |
| Anostomidae | *Schizodon* | *Schizodon fasciatus* | 0 | 6 | 2 |
| Anostomidae | *Schizodon* | *Schizodon* sp. | 6 | 0 | 0 |
| Auchenipteridae | *Auchenipterus* | *Auchenipterus ambyiacus* | 0 | 0 | 2 |
| Auchenipteridae | *Auchenipterus* | *Auchenipterus*<br>*ambyiacus/Auchenipterus nuchalis* | 0 | 2 | 0 |
| Auchenipteridae | *Auchenipterus* | *Auchenipterus* sp. | 2 | 0 | 0 |
| Auchenipteridae | *Trachelyopterus* | *Trachelyopterus galeatus* | 3 | 1 | 0 |
| Auchenipteridae | *Trachelyopterus* | *Trachelyopterus porosus* | 0 | 0 | 1 |
| Belonidae | *Potamorhaphis* | *Potamorhaphis* cf. *eigenmanni* | 0 | 0 | 1 |
| Cetopsidae | *Cetopsis* | *Cetopsis coecutiens* | 0 | 0 | 1 |
| Characidae | *Aphyocharax* | *Aphyocharax avaray* | 0 | 0 | 6 |
| Characidae | *Aphyocharax* | *Aphyocharax* sp. | 0 | 2 | 0 |
| Characidae | *Aphyocheirodon* | *Aphyocheirodon* sp. | 0 | 6 | 0 |
| Characidae | *Astyanax* | *Astyanax bimaculatus* | 6 | 9 | 5 |
| Characidae | *Brachyhalcinus* | *Brachyhalcinus* aff. *nummus* | 0 | 0 | 1 |
| Characidae | *Brachyhalcinus* | *Brachyhalcinus copei* | 0 | 0 | 2 |
| Characidae | *Charax* | *Charax gibbosus* | 0 | 2 | 4 |
| Characidae | *Charax* | *Charax pauciradiatus* | 0 | 0 | 1 |
| Characidae | *Compsura* | *Compsura* sp. | 2 | 0 | 0 |
| Characidae | *Ctenobrycon* | *Ctenobrycon hauxwellianus* | 9 | 0 | 7 |
| Characidae | *Cynopotamus* | *Cynopotamus* sp. | 3 | 0 | 0 |
| Characidae | *Galeocharax* | *Galeocharax gulo* | 0 | 0 | 1 |
| Characidae | *Gymnocorymbus* | *Gymnocorymbus ternetzi* | 8 | 0 | 1 |
| Characidae | *Gymnocorymbus* | *Gymnocorymbus thayeri* | 0 | 0 | 1 |
| Characidae | *Hemigrammus* | *Hemigrammus* sp. | 2 | 0 | 3 |
| Characidae | *Jupiaba* | *Jupiaba anteroides* | 0 | 0 | 1 |
| Characidae | *Knodus* | *Knodus* sp. 1 | 0 | 0 | 1 |
| Characidae | *Knodus* | *Knodus* sp. 2 | 0 | 0 | 1 |
| Characidae | *Knodus* | *Knodus* sp. 3 | 0 | 0 | 1 |
| Characidae | *Knodus* | *Knodus* sp. 4 | 0 | 0 | 1 |
| Characidae | *Knodus* | *Knodus* sp. 5 | 0 | 0 | 3 |
| Characidae | *Knodus* | *Knodus* sp. 6 | 0 | 0 | 4 |
| Characidae | *Moenkhausia* | *Moenkhausia bonita* | 0 | 0 | 1 |
| Characidae | *Moenkhausia* | *Moenkhausia lepidura* | 0 | 0 | 1 |
| Characidae | *Moenkhausia* | *Moenkhausia madeirae* | 0 | 0 | 7 |
| Characidae | *Moenkhausia* | *Moenkhausia oligolepis* | 0 | 0 | 1 |
| Characidae | *Moenkhausia* | *Moenkhausia sanctaefilomenae* | 2 | 0 | 0 |
| Characidae | *Moenkhausia* | *Moenkhausia* sp. | 2 | 6 | 0 |
| Characidae | *Phenacogaster* | *Phenacogaster* sp. | 4 | 0 | 0 |
| Characidae | *Poptella* | *Poptella compressa* | 1 | 0 | 0 |
| Characidae | *Roeboides* | *Roeboides afinnis* | 0 | 0 | 1 |
| Characidae | *Roeboides* | *Roeboides biserialis* | 0 | 0 | 1 |
| Characidae | *Roeboides* | *Roeboides descalsvadensis* | 0 | 3 | 0 |
| Characidae | *Roeboides* | *Roeboides* sp. | 5 | 2 | 0 |
| Characidae | *Serrapinnus* | *Serrapinnus piaba* | 7 | 0 | 0 |
| Characidae | *Serrapinnus* | *Serrapinnus* sp. 1 | 0 | 0 | 6 |
| Characidae | *Serrapinnus* | *Serrapinnus* sp. 2 | 0 | 0 | 1 |
| Characidae | *Tetragonopterus* | *Tetragonopterus argenteus* | 0 | 2 | 1 |
| Characidae | *Tyttobrycon* | *Tyttobrycon* sp. | 0 | 0 | 1 |
| Cichlidae | *Aequidens* | *Aequidens tetramerus* | 0 | 0 | 2 |
| Cichlidae | *Apistogramma* | *Apistogramma* sp. | 0 | 0 | 3 |
| Cichlidae | *Bujurquina* | *Bujurquina eurhinus* | 0 | 0 | 2 |
| Cichlidae | *Cichlasoma* | *Cichlasoma bimaculatum* | 0 | 10 | 0 |
| Cichlidae | *Cichlasoma* | *Cichlasoma boliviense* | 0 | 0 | 5 |
| Cichlidae | *Cichlasoma* | *Cichlasoma* sp. | 6 | 1 | 0 |
| Cichlidae | *Mesonauta* | *Mesonauta festivus* | 0 | 2 | 2 |
| Cichlidae | *Satanoperca* | *Satanoperca jurupari* | 0 | 0 | 5 |
| Cichlidae | *Satanoperca* | *Satanoperca* sp. | 0 | 5 | 0 |
| Cichlidae | *Saxatilia* | *Saxatilia lepidota* | 10 | 0 | 0 |
| Cichlidae | *Saxatilia* | *Saxatilia semicincta* | 0 | 0 | 5 |
| Cichlidae | *Saxatilia* | *Saxatilia* sp. | 0 | 9 | 0 |
| Crenuchidae | *Characidium* | *Characidium* sp. | 0 | 0 | 1 |
| Curimatidae | *Curimatella* | *Curimatella meyeri* | 0 | 0 | 1 |
| Curimatidae | *Curimatella* | *Curimatella* sp. | 0 | 2 | 0 |
| Curimatidae | *Cyphocharax* | *Cyphocharax* sp. | 6 | 5 | 0 |
| Curimatidae | *Cyphocharax* | *Cyphocharax spiluroopsis* | 0 | 0 | 3 |
| Curimatidae | *Potamorhina* | *Potamorhina altamazonica* | 0 | 8 | 5 |
| Curimatidae | *Psectrogaster* | *Psectrogaster rutiloides* | 0 | 2 | 2 |
| Curimatidae | *Steindachnerina* | *Steindachnerina aff. dobula* | 0 | 0 | 4 |
| Curimatidae | *Steindachnerina* | *Steindachnerina guentheri* | 0 | 2 | 3 |
| Curimatidae | *Steindachnerina* | *Steindachnerina* sp. | 0 | 0 | 1 |
| Cynodontidae | *Cynodon* | *Cynodon gibbus* | 0 | 0 | 1 |
| Cynodontidae | *Cynodon* | *Cynodon* sp. | 0 | 1 | 0 |
| Cynodontidae | *Hydrolycus* | *Hydrolycus scomberoides* | 1 | 0 | 0 |
| Erythrinidae | *Hoplerythrinus* | *Hoplerythrinus unitaeniatus* | 0 | 2 | 2 |
| Erythrinidae | *Hoplias* | *Hoplias malabaricus* | 9 | 8 | 9 |
| Gymnotidae | *Gymnotus* | *Gymnotus carapo* | 6 | 4 | 0 |
| Hemiodontidae | *Anodus* | *Anodus elongatus* | 0 | 0 | 1 |
| Heptapteridae | *Pimelodella* | *Pimelodella* sp. | 1 | 0 | 0 |
| Hypopomidae | *Brachyhypopomus* | *Brachyhypopomus pinnicaudatus* | 0 | 1 | 0 |
| Hypopomidae | *Brachyhypopomus* | *Brachyhypopomus* sp. | 3 | 0 | 0 |
| Iguanodectidae | *Bryconops* | *Bryconops* sp. | 1 | 0 | 0 |
| Iguanodectidae | *Bryconops* | *Bryconops melanurus* | 0 | 0 | 1 |
| Loricariidae | *Ancistrus* | *Ancistrus* sp. | 5 | 4 | 0 |
| Loricariidae | *Hypoptopoma* | *Hypoptopoma* cf. *bianale* | 0 | 0 | 1 |
| Loricariidae | *Hypoptopoma* | *Hypoptopoma gulare* | 0 | 4 | 1 |
| Loricariidae | *Hypoptopoma* | *Hypoptopoma* sp. | 5 | 0 | 0 |
| Loricariidae | *Hypostomus* | *Hypostomus* sp. | 8 | 7 | 0 |
| Loricariidae | *Hypostomus* | *Hypostomus* sp.1 | 0 | 0 | 5 |
| Loricariidae | *Hypostomus* | *Hypostomus* sp.2 | 0 | 0 | 2 |
| Loricariidae | *Loricariichthys* | *Loricariichthys platymetopon* | 7 | 7 | 4 |
| Loricariidae | *Loricariichthys* | *Loricariichthys* sp. | 0 | 0 | 1 |
| Loricariidae | *Pterygoplichthys* | *Pterygoplichthys disjunctivus* | 0 | 0 | 1 |
| Pimelodidae | *Hemisorubim* | *Hemisorubim platyrhynchos* | 1 | 0 | 0 |
| Pimelodidae | *Hypophthalmus* | *Hypophthalmus edentatus* | 2 | 0 | 1 |
| Pimelodidae | *Hypophthalmus* | *Hypophthalmus* sp. | 0 | 2 | 0 |
| Pimelodidae | *Pimelodus* | *Pimelodus* sp. | 4 | 0 | 0 |
| Pimelodidae | *Pimelodus* | *Pimelodus tetramerus* | 0 | 2 | 0 |
| Pimelodidae | *Pseudoplatystoma* | *Pseudoplatystoma* sp. | 1 | 0 | 0 |
| Pimelodidae | *Pseudoplatystoma* | *Pseudoplatystoma tigrinum* | 3 | 0 | 0 |
| Pimelodidae | *Sorubim* | *Sorubim lima* | 2 | 1 | 1 |
| Prochilodontidae | *Prochilodus* | *Prochilodus lineatus* | 8 | 0 | 0 |
| Prochilodontidae | *Prochilodus* | *Prochilodus nigricans* | 0 | 5 | 4 |
| Serrasalminae | *Mylossoma* | *Mylossoma albiscopum* | 0 | 1 | 1 |
| Serrasalminae | *Serrasalmus* | *Serrasalmus maculatus* | 0 | 0 | 5 |
| Serrasalminae | *Serrasalmus* | *Serrasalmus rhombeus* | 0 | 3 | 3 |
| Serrasalminae | *Serrasalmus* | *Serrasalmus* sp. | 6 | 0 | 0 |
| Serrasalminae | *Serrasalmus* | *Serrasalmus spilopleura* | 0 | 0 | 2 |
| Sternopygidae | *Eigenmannia* | *Eigenmannia* cf. *macrops* | 0 | 0 | 1 |
| Sternopygidae | *Eigenmannia* | *Eigenmannia* gr. *trilineata* | 0 | 1 | 0 |
| Sternopygidae | *Eigenmannia* | *Eigenmannia limbata* | 0 | 1 | 0 |
| Sternopygidae | *Eigenmannia* | *Eigenmannia* sp. | 3 | 2 | 1 |
| Sternopygidae | *Eigenmannia* | *Eigenmannia virescens* | 0 | 0 | 1 |
| Sternopygidae | *Sternopygus* | *Sternopygus macrurus* | 8 | 6 | 2 |
| Synbranchidae | *Synbranchus* | *Synbranchus marmoratus* | 1 | 0 | 1 |
| Triportheidae | *Triportheus* | *Triportheus angulatus* | 0 | 0 | 4 |
| Triportheidae | *Triportheus* | *Triportheus rotundatus* | 0 | 0 | 1 |
| Triportheidae | *Triportheus* | *Triportheus* sp. | 0 | 6 | 0 |

When considering the sites surveyed independently, we could identify more morphospecies when using the eDNA approach with both markers combined than with traditional methods (TM) only, with the exception of Chambira FP#1, a flooded abandoned mining pond, where TM identified three additional morphospecies in contrast to the eDNA method (Figure 5). It is important to note that at this site, and others, many MOTUs were not mapped at the morphospecies level (Table 2). Interestingly, it would appear that greater species richness was obtained by combining eDNA and traditional methods (Figure 5); however, given that the identity of the fish obtained with traditional methods was not verified using 12S and COI markers, it cannot be ruled out that some taxonomically close morphospecies detected by either method are actually the same.

**Figure 5.**
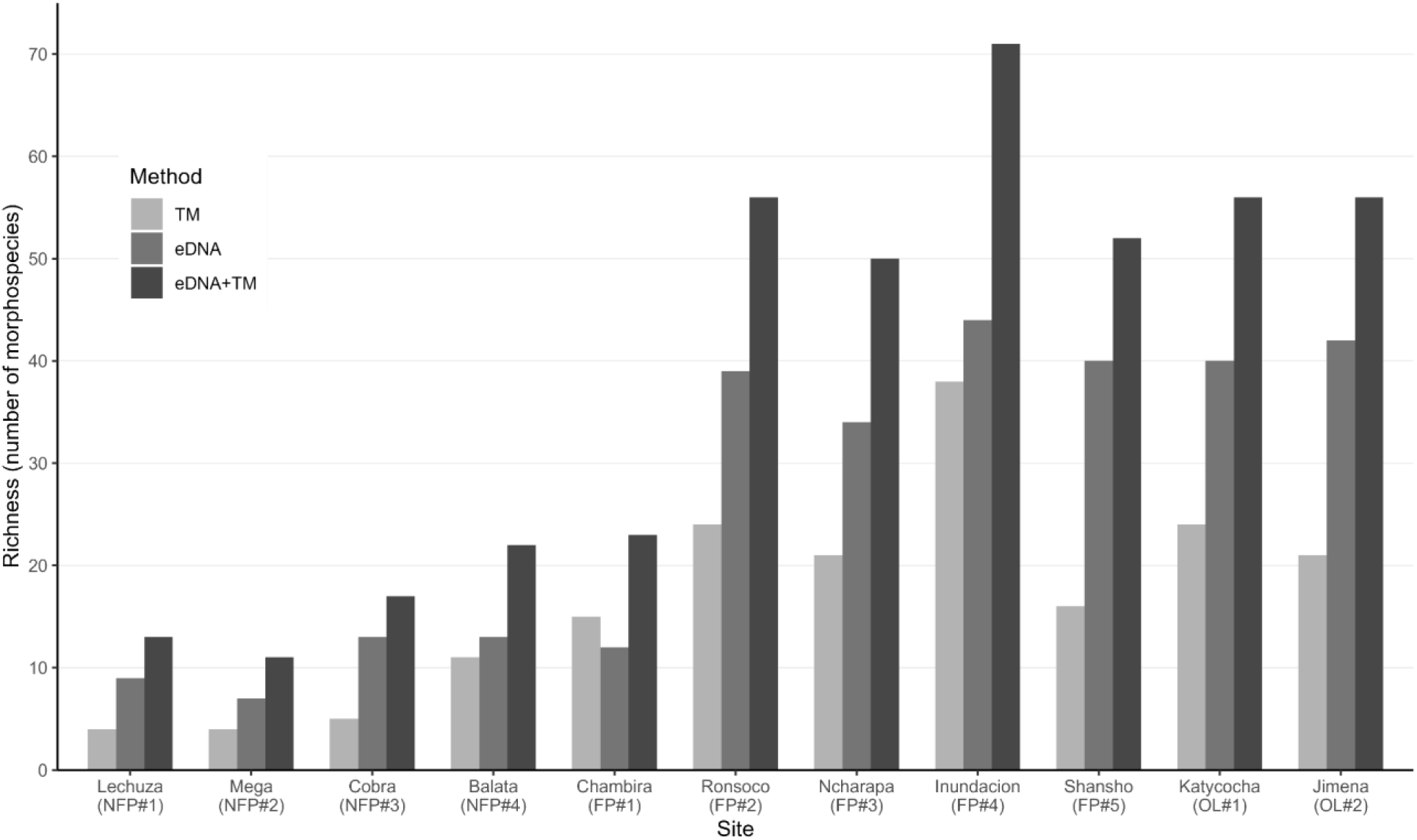
Total number of morphospecies registered at each site using only traditional methods (TM), using eDNA - compiling results obtained with the12S and COI markers (eDNA) −, and using traditional methods and eDNA combined (TM + eDNA). MOTUs not mapped at morphospecies level are not included in the graph. NFP: Non-flooded [abandoned mining] ponds. FP: Flooded [abandoned mining] ponds. OL: [pristine] Oxbow lakes.

At the family level, both approaches identified 5 families not detected with the other approach (Figure 6). At the genera level, the eDNA method identified 14 genera not found with the traditional method, whereas the traditional approach found 13 genera not registered by eDNA (Figure 6). Finally, at the morphospecies level, the eDNA method identified 47 morphospecies not found by the traditional methods, whereas the TM identified 53 morphospecies not registered by eDNA (Figure 6). Both methods identified 15 shared families, 40 shared genera, and 23 shared morphospecies (Figure 6).

**Figure 6.**
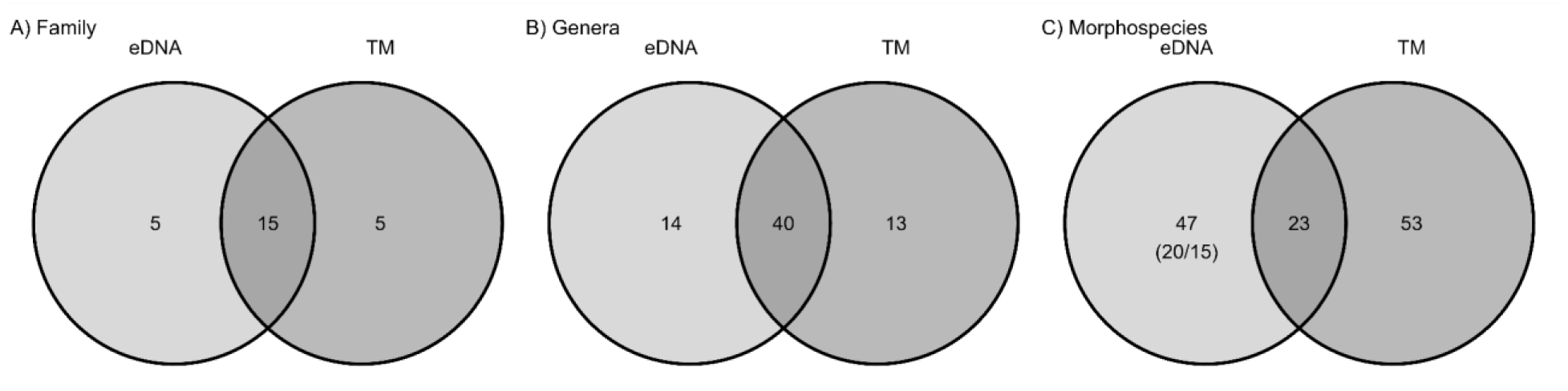
Venn diagram showing the number of families, genera and morphospecies shared by both methods: eDNA and Traditional methods (TM). Numbers in parenthesis indicate MOTUS not mapped at morphospecies level by the 12S/COI markers respectively.

Regarding commercial species, we identified more diversity with eDNA than with traditional approaches (Table 3). With the 12S marker we registered seven species: *Hydrolycus scomberoides* (common name, chambira or cachorro), *Hemisorubim platyrhynchos* (common name, pico pato or toa), *Pseudoplatystoma tigrinum* (common name, puma zungaro or tiger sorubim), *Pseudoplatystoma* sp. (common name, doncella or barred sorubim), *Prochilodus lineatus* (common name, bocachico or curimata), *Leporinus* sp., and *Schizodon* sp. (common name, both as lisa or araçu). With the COI marker we registered 10 species: *Schizodon fasciatus*, *Leporinus* aff. *friderici*, *Leporinus* sp. (common name, both as lisa or araçu), *Megaleporinus trifasciatus* (common name, lisa or araçu), *Potamorhina altamazonica* (common name, yahuarachi or llambina), *Psectrogaster rutiloides* (common name, chio-chio or branquinha), *Prochilodus nigricans* (common name, bocachico or curimata), *Mylossoma albiscopum* (common name, palometa or pacupeba), *Pimelodus tetramerus* (common name, bagre-cunchi or bloch’s catfish) and *Sorubim lima* (common name, pico pato - shiripira or duckbill catfish).

With the eDNA method we were able to identified species that were difficult to capture by traditional methods such as the electric fishes *Gymnotus carapo*, *Brachyhypopomus* sp, *Brachyhypopomus pinnicaudatus*, and *Eigenmannia limbata* from the Order Gymnotiformes which are usually found around trees, and beneath branches and leaf litter from macrophytes and riparian vegetation in the ponds.

Finally, using both methods, we found that some species were more prevalent in the surveyed landscape, being found in almost all sites. These include *Saxatilia lepidota*-formerly *Crenicichla lepidota-*, *Cichlasoma bimaculatum*, *Leporinus* sp., *Ctenobrycon hauxwellianus*, *Astyanax bimaculatus*, *Saxatilia* sp., and the species *Hoplias malabaricus*, from the Erythrinidae family, which was found at all sites surveyed using either method and marker (supplementary information 8).

### Effect of ecological context on fish diversity

We investigated the effect of flood pulse, year of abandonment, and pond dimension on richness (as number of morphospecies) using ANOVA. The degree of flood pulse had a statistically significant effect on fish richness (p-value < 0.05), whereas the year of abandonment (*p-value* = 0.804) and surface area (*p-value* = 0.183) did not. We identified a total of 123 morphospecies across all sites using traditional and eDNA monitoring combined (Table 3), of which 32 were found in non-flooded ponds (NFP), 101 in flooded ponds (FP), and 76 in oxbow lakes (OL). NFPs exhibit lower species richness (16 ± 5 SD per site) compared to both FP (50 ± 17 SD per site) and OL (56 ± 0 SD per site) (Figure 5 and Table 2). Overall, our results indicate that seasonal flood pulse plays a crucial role in influencing aquatic communities and species accumulation, as suggested in Araujo-Flores et al. (2021).

Following the ANOVA results, we decided to further test the effect of flood pulse on fish community composition using a PERMANOVA test. We found statistically significant differences in fish species composition among ponds affected by different flood pulse conditions (p-value < 0.005). When plotting the Jaccard dissimilarity index in a principal coordinate analysis (PCoA) we could see that the fish community composition among OL and FP had high similarity - excluding FP#1-in contrast to the non flooded ponds (Figure 7). The NFP#1 displayed a distinct ordination pattern among NFPs, possibly due to its central location within La Pampa, being the least connected to streams or forest (see Figure 1).

**Figure 7.**
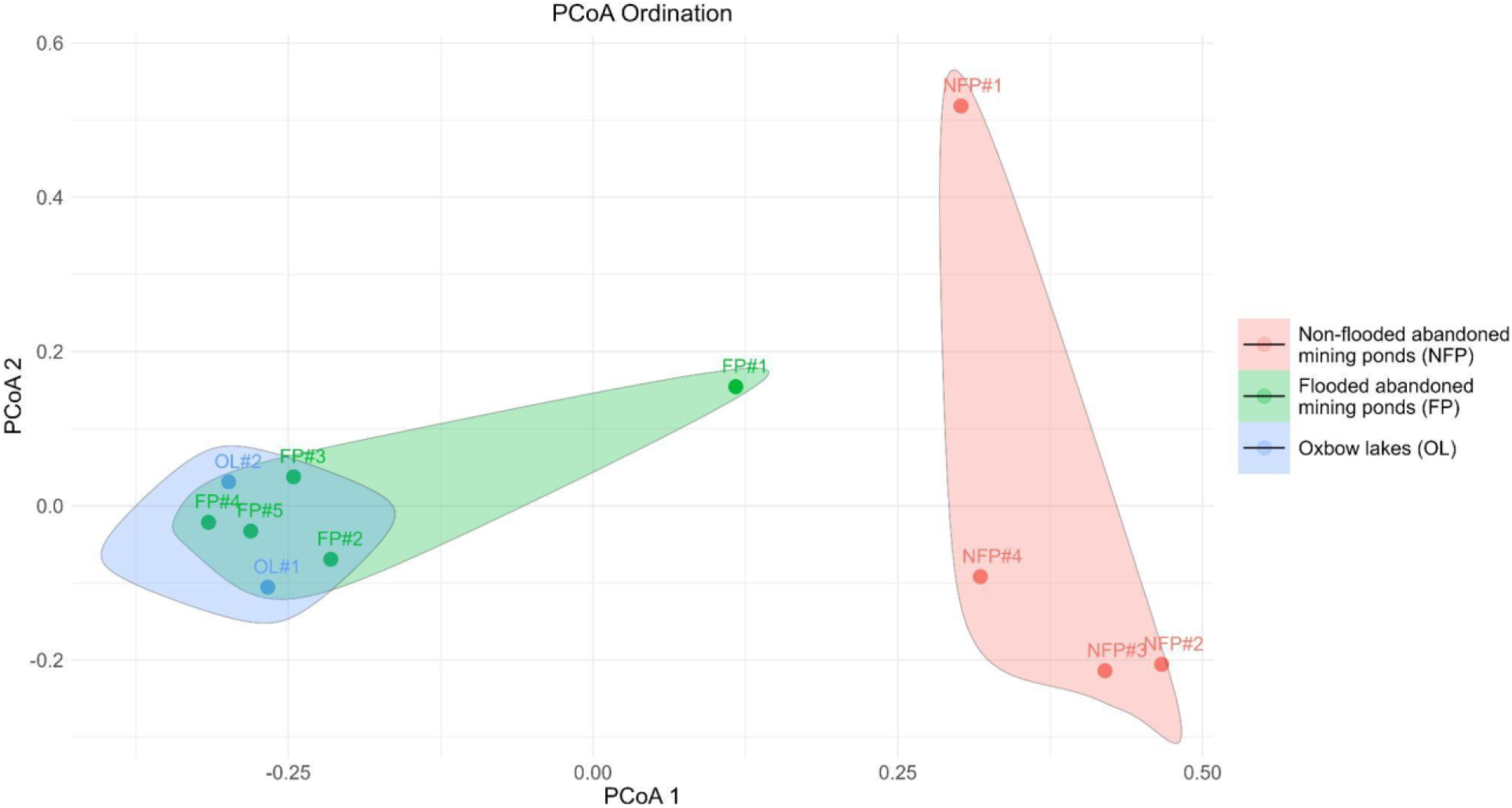
PCoA Ordination using Jaccard dissimilarity index.

The NFPs, located in La Pampa, had similar fish composition among them, with a predominance of the Cichlidae and Erythrinidae family (Figure 4). The genus *Saxatilia* was predominant in the NFP ecosystems, with a greater proportion of reads per sample (see supplementary information 2 and 3), and a higher abundance found with traditional methods (see supplementary information 5). While the FP and OL share more genera and morphospecies between them, compared to NFP and OL, or between NFP and FP, when looked on proportion level, the NFP shared 62.5% of their diversity with both conditions, while FP and OL shared only 19.8% and 26.3%, respectively (Figure 8). This is due to an important proportion of morphospecies found in NFPs are species that probably can survive in almost all mining ponds and oxbow lakes (Figure 8). Interestingly, the FP, which are influenced by the Madre de Dios and Malinowski River (see Table 1), have a higher richness of fish when compared to the OL, influenced by the La Torre River, a smaller hierarchy river (Figure 5).

**Figure 8.**
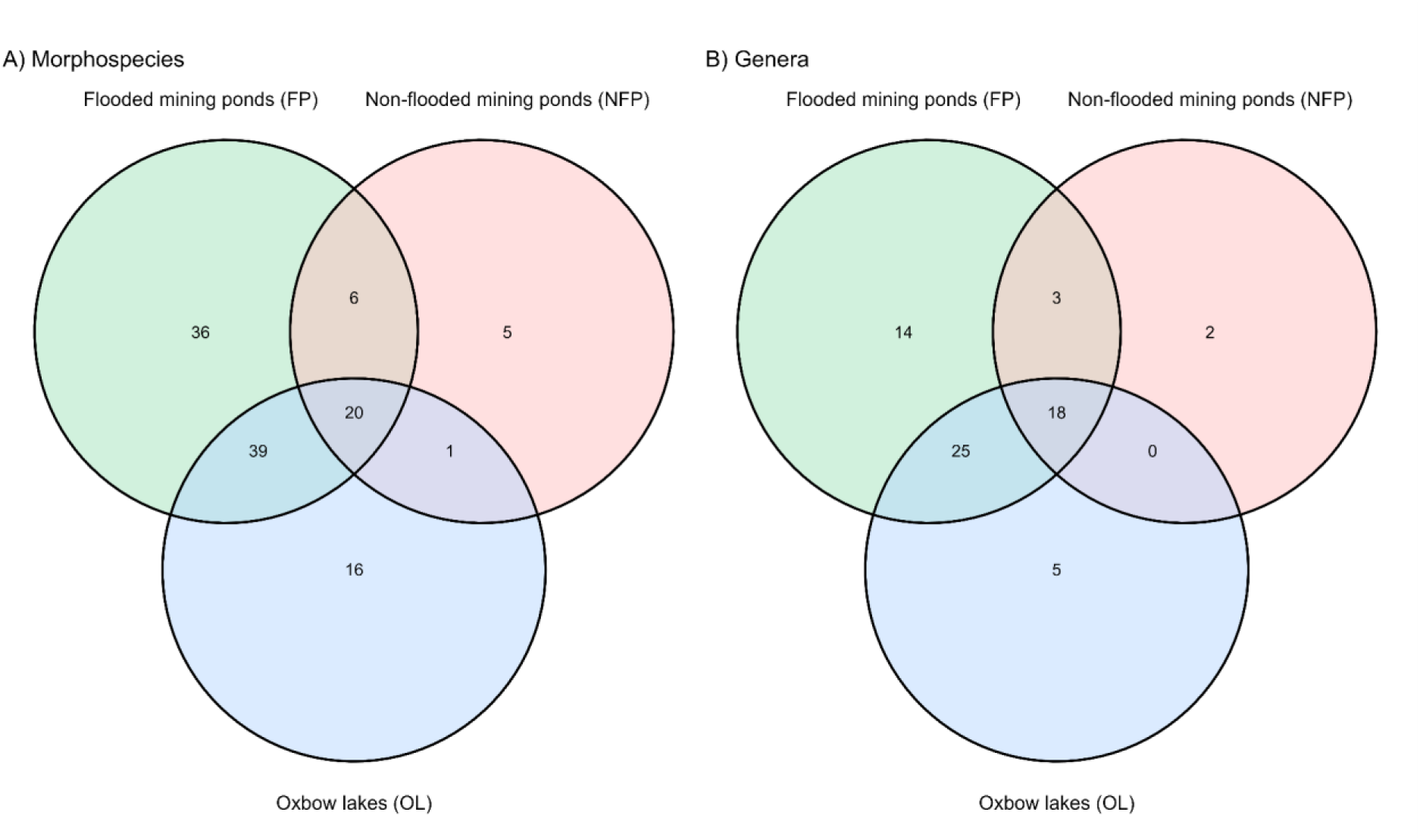
Venn-diagram showing the number of morphospecies and genera shared by the three different conditions (NFP, FP, OL), considering both approaches eDNA and TM combined.

## Discussion

In this study, we assessed fish diversity (i.e., richness) and community composition of abandoned mining ponds using both eDNA and traditional monitoring to understand factors that could be influencing the recolonization dynamics of post-ASGM landscapes within the Madre de Dios region, to evaluate possible recovery strategies. We studied nine mining ponds with different degree of pulse flood connectivity with extant waterways, classified as flooded mining ponds (FP), located in a seasonal flooded ecosystem, and non-flooded mining ponds, located in a *terra firme* or non-flooded ecosystem (NFP), with and without connection to lotic systems, with different age since the abandonment and different dimensions, and two nearby unmined oxbow lakes (OL) used as reference (see Table 1). Overall, our results highlight the value of eDNA as a cost-efficient tool for rapid biodiversity monitoring (Andres et al., 2023; Pukk et al., 2021), especially in remote and hard to access areas (Timana-Mendoza et al., 2024).

### eDNA fish primers performance

We used two molecular markers for detecting fish diversity: the COI marker (Mariac et al., 2022) and the 12S marker (Riaz et al., 2011 in Kelly et al., 2014). The COI gene is a well-established marker for assessing animal diversity, particularly supported by its extensive database (Collins et al., 2019; Teixeira et al., 2023). While its high interspecific variability makes it suitable for differentiating closely related species, it requires highly degenerate primers, which limits its effectiveness in broader biodiversity detection (Kumar et al., 2022; Teixeira et al., 2023; Zhang et al., 2020). In contrast, the 12S gene is widely utilized in metabarcoding due to its conserved regions that facilitate primer design and its variable regions that allow genus- or species-level resolution, enhancing the detection of fish taxa (Jackman et al., 2021; Kumar et al., 2022).

In our study, both molecular markers successfully characterized fish diversity, detecting a similar number of morphospecies across all sites, despite some differences. For instance, the 12S marker outperformed the COI marker by identifying more MOTUs per location (Figure 2). However, the COI marker identified 10 additional MOTUs at the species level compared to the 12S marker (supplementary information 3), indicating higher taxonomic resolution, probably due to the availability of a more extensive database (Collins et al., 2019; Jackman et al., 2021; Milan et al., 2020, among others).

Nevertheless, the COI marker exhibited limitations, including four instances where a single identified MOTU was assigned to two different species with 100% sequence similarity, reflecting its reduced ability to differentiate between these species (Zhang et al., 2020). Furthermore, the COI marker recorded 566 MOTUs, representing a significant proportion of the reads (1,562,389 reads), that were unassigned or corresponded to non-target MOTUs, indicating poor target specificity (Collins et al., 2019; Kumar et al., 2022; Zhang et al., 2020). In contrast, the 12S marker had only 27 unassigned MOTUs (358,211 reads). Despite these differences, each marker provided unique fish diversity information (Figure 3 and Figure 4), highlighting the importance of employing multiple primer sets to improve taxonomic coverage (Garcia-Machado et al., 2023; Munian et al., 2024), particularly in megadiverse regions (Jackman et al., 2021; Zhang et al., 2020).

### eDNA and traditional approach monitoring

When comparing the two monitoring approaches used in the study, eDNA identified a greater number of morphospecies at nearly all sites (Figure 5), highlighting its advantages for detecting fish diversity (Munian et al., 2024; Pont et al., 2023; Thomsen et al., 2016; Yao et al., 2022). Additionally, it required fewer resources than traditional methods, as noted by Timana-Mendoza et al. (2024). eDNA also detected the presence of some species that are usually not recorded using traditional methods due to their elusive habits or the extensive fishing effort required, such as *Trachelyopterus galeatus, Pseudoplatystoma tigrinum, Sternopygus macrurus, Gymnotus carapo, Saxatilia* spp., *Hypoptopoma* spp., *Ancistrus* spp., and *Brachypopomus* spp. (Table 3).

Nevertheless, species detection via eDNA in Amazon aquatic environments remains challenging due to limitations in taxonomic resolution, which can prevent some MOTUs from being identified at the species level or result in misannotations in genomic databases (Beng & Corlett, 2020; Collins et al., 2019; Mariac et al., 2022; Yao et al., 2022). For instance, in our study, the 12S marker identified *Prochilodus lineatus*, a species not found in the Madre de Dios River basin or the Beni-Madeira River system (Carvajal-Vallejos & Zeballos-Fernández, 2011; Queiroz et al., 2013; Ortega et al., 2012). In contrast, *Prochilodus nigricans* (commonly known as bocachico or curimata), an important species for subsistence fishing throughout the Amazon (García-Dávila et al., 2018; Goulding et al., 2003; Silvano, 2020), was identified using the COI marker and traditional methods, and may represent the same species.

Similarly, the COI marker identified *Satanoperca* sp., while traditional methods identified *Satanoperca jurupari*, which could be an unrecorded species in genomic databases or a potential new species (Willis et al., 2012). Additionally, the genus *Moenkhausia* was represented by one morphospecies with the COI marker (*Moenkhausia* sp.), two morphospecies with the 12S marker (*Moenkhausia* sp. and *Moenkhausia sanctaefilomenae*), and four species through traditional methods (*Moenkhausia bonita*, *Moenkhausia lepidura*, *Moenkhausia madeirae*, and *Moenkhausia oligolepis*). Notably, *Moenkhausia sanctaefilomenae*, identified with the 12S marker, is not recorded in Peru, suggesting a possible misidentification in genomic databases and potentially corresponding to one of the species identified using traditional methods. Therefore, as previously reported in Neotropical freshwater ecosystems, the limited reference database for fish constrains our results, emphasizing the need for continuous sequencing efforts to expand reference sequence databases and to correctly annotate species (Coutant et al., 2023; Hilário et al., 2023; Milan et al., 2020).

Taxonomic identification of fish using traditional methods is also limited by their time-consuming nature and often controversial results, especially for morphologically challenging ichthyofauna or species lacking reliable taxonomic keys (e.g., *Brachyopomus*, *Eigenmannia*, *Loricariichthys*, *Hypostomus*, *Serrapinnus*, *Moenkhausia*, *Knodus*, and others) (Araújo-Flores, 2016; van der Sleen & Albert, 2017). For example, traditional methods recorded 6 different species from the *Knodus* genera, but no taxonomic keys are available to distinguish them. These methods also detected species absent in eDNA samples, possibly due to their position in the water column (Rehill et al., 2024; Rourke et al., 2022), small size, or low frequency (e.g., *Aequidens tetramerus*, *Anodus elongatus*, *Cetopsis coecutiens*, *Pterygoplichthys disjunctivus*, *Potamorrhaphis* cf. *eigenmanni*, *Bryconops melanurus*, *Apistogramma* spp.) (Galvis et al., 2006; Van der Sleen & Albert 2017). All of these mentioned genera and species were commonly reported using traditional fishing methods yet remained undetected by the 12S and COI markers (Table 3).

Multiple studies have highlighted the effectiveness of molecular techniques for detecting invasive fish species, particularly in temperate regions (Duarte et al., 2023; Nynatten et al., 2023; Takahashi et al., 2023). Despite this, our study did not detect the presence of invasive species previously reported in our river basin, such as *Arapaima* (Paiche or Pirarucu), *Oreochromis* (Tilapia) in the floodplain, and *Oncorhynchus* (Trout or Trucha) in the Andean foothills (Goulding et al., 2014; Araújo-Flores, 2016; Pelicice & Rodrigues, 2021; Catâneo et al., 2022). The development of molecular techniques for diversity monitoring in these new wetlands represents a promising opportunity to manage this vast and ever-changing portion of the territory, transformed by mining activities.

Fish biodiversity monitoring using traditional methods in the study area can compromise fieldwork safety, especially with techniques like electrofishing, dip nets, seines, and gillnets, which require extended time at sites. In our study area, there is a reported social instability due to mining activities (Dethier et al., 2023b), which has changed the dynamics of settlements in the mining sector of La Pampa, that currently prevent us from accessing locations that were previously considered safe and had the potential to host long-term ecological studies, as reported in our four ASGM ponds: Lechuza NFP#1, Mega NFP#2, Cobra NFP#3, and Balata NFP#4 (Araujo-Flores personal communication). Finally, the eDNA approach was able to identify other vertebrate taxa, 10 and 29 MOTUs with the COI and 12S marker, respectively, within the classes Mammalia, Reptilia, Amphibians, and Aves (supplementary information 6 and 7). Notably, *Lontra longicaudis* (‘river otter’, Near Threatened), *Podocnemis unifilis* (‘yellow-headed sideneck turtle’ or ‘taricaya’, Vulnerable), *Tapirus terrestris* (‘tapir’, Vulnerable), and *Alouatta seniculus* (‘red howler monkeys’, Least Concern) were found in mining ponds as well as pristine lakes. The latter species was detected in Mega NFP#2, an area with significant mining activity, raising safety concerns. These findings underscore the importance of management strategies to protect vulnerable and near-threatened species within these aquatic ecosystems.

### Species richness and fish communities

Our study revealed that seasonal flooding plays a crucial role in fish recolonization of mining ponds, as previously suggested (Araujo-Flores et al., 2021). We found significant differences in fish species richness (Figure 5) and community composition (Figure 7) when analyzing all three conditions of study sites with different degrees of connectivity to lotic systems (i.e., FP, NFP, and OL), in contrast to the abandonment time and the dimensions of the pond or lake. We found that the FP (5 mining ponds) exhibited the highest number of morphospecies and MOTUs, followed by OL (2 lakes), and NFP (4 mining ponds). Although the abandonment time of the mining ponds should allow more time for ecosystem restoration or recolonization, the lack of connection between NFPs and lotic systems prevents these ponds from benefiting from incoming aquatic communities, nutrients, and resources (Pander et al., 2018). For example, Lechuza NFP#1 and Mega NFP#2, located in La Pampa and not connected to lotic systems or forests, had the lowest richness among the study sites despite being the largest mining ponds and having been abandoned since 2011. Similarly, the Cobra NFP#3 and Balata NFP#4 sites, also located in La Pampa, though slightly influenced by a small stream from the Malinowski River, also presented relatively low diversity (Table 2) due to the lack of connection with a river main channel. Moreover, the similarity in species composition between Chambira FP#1 and NFP (Figure 7 and supplementary information 1) suggests less restoration in this mining pond, which was also observed during field sample collection. Furthermore, the disappearance of migratory fish, many of which are of commercial interest, from the mining ponds of the Malinowski River (FP#1 and FP#2), a tributary heavily impacted by mining, suggests the deterioration of these ecosystems due to poor connectivity with the rest of the basin, hindering access to these areas for spawning (Arana & Chang, 2005; Duponchelle et al., 2021).

Notably, families such as Cichlidae, Characidae, Erythrinidae, and Curimatidae were present across almost all sites, which could indicate their role in recolonization and adaptation to various conditions. Furthermore, the majority of the taxa found in NFP were present in all three conditions (Figure 8), indicating that the species in this ecosystem could be the first to recolonize environments with limited nutrients and resources, such as the genera Saxatilia and Cichlasoma, which are dominant in NFP sites.

Finally, FP exhibit even higher richness than OL (Table 2), such as the Inundacion FP#4 and the Ronsoco FP#2 sites, influenced by the Madre de Dios River and the Malinowski River, respectively. This suggests potential ecosystem recovery, offering a hopeful perspective for post-mining restoration efforts. The lower richness in OL may result from their connection to the smaller La Torre River, which provides fewer nutrients and species compared to the Madre de Dios River, known for its critical role in seasonal flooding (Fernandes et al., 2014; Strahler, 1952; Timana-Mendoza et al., 2024). Our study highlights the need for more pristine lakes to improve our understanding of landscape ecology (Terborgh et al., 2018).

## Conclusions

Our study demonstrates the efficacy of eDNA as a powerful, cost-efficient tool for monitoring fish diversity in post-ASGM landscapes, complementing traditional monitoring methods. Both 12S and COI molecular markers provided valuable insights, with the 12S marker outperforming in taxonomic coverage and identification of unique taxa, while the COI marker offered higher species-level resolution due to a more extensive reference database. Therefore, using multiple molecular markers provides a more comprehensive understanding of species richness in both mining ponds and oxbow lakes. While eDNA identified a higher number of taxa, traditional methods still detected unique species not captured by molecular techniques. Our results emphasize the need for continued sequencing efforts to expand genomic reference databases for Amazonian fishes in order to accurately identify species in post-mining ecosystems, including threatened species, to better inform conservation and restoration strategies. Notably, we found that the connectivity of the mining ponds to lotic systems plays a significant role in species richness and community composition in mining ponds. Flooded mining ponds exhibited species richness and assemblages almost comparable to those of unimpacted oxbow lakes, suggesting potential ecosystem recovery, which can be a hopeful perspective for post-mining restoration efforts. In contrast, mining ponds disconnected from lotic systems showed drastically reduced biodiversity, despite being abandoned for more years and with a greater pond dimension. However, we observed that some mammals use these areas, highlighting their importance for certain species and the need for ecosystem management. Overall, our findings provide critical insights into the recovery and biodiversity of post-ASGM ponds and support the integration of both eDNA and traditional methods for monitoring freshwater ecosystems in the Amazon basin.

## Acknowledgments

This work was supported by the generous support of the American people through the United States Agency for International Develepment (USAID; Grant N°. 72052721CA005) bestowed to the Center of Amazonian Scientific Innovation (CINCIA); and the support of the Peruvian National Council of Science, Technology and Technological Innovation, under the project PE501082020-2023. The contents of this paper do not necessarily reflect the views of USAID or the United States Government. The authors would also like to thank the Servicio Nacional de Áreas Protegidas por el Estado (SERNANP) and Ejército del Perú (EP) for the logistical support that made possible the development of this study. This study was carried on under permit: RESOLUCIÓN DIRECTORAL N° 00476-2022-PRODUCE/DGPCHDI, RESOLUCIÓN DIRECTORAL N° 00812-2023-PRODUCE/DGPCHDI and RESOLUCION JEFATURAL DE LA RESERVA NACIONAL TAMBOPATA N° 09-2022-SERNANP-JEF. The authors are also thankful to their field assistants for their strong commitment and work in difficult field conditions, Dr. Junior Chuctaya for his revision of fish species present in Peru, BSc. Renzo Rivera for some fish taxonomic identification, and the LIDAE Program from CINCIA for the floodplain status and the abandonment year identification.

## Data Accessibility and Benefit-Sharing

The raw data used for this study is presented as supplementary information

## Author Contributions

CTM: experimental design, fieldwork, analysis and interpretation of the data, manuscript preparation, manuscript revision

ARC: laboratory work, manuscript preparation

PV: interpretation of the data, manuscript revision

RB: taxonomic identification assistance, manuscript revision

MCSM: experimental design, analysis and interpretation of the data, manuscript preparation, manuscript revision

JMAF: secured funding, experimental design, fieldwork, interpretation of the data, manuscript revision

MS: secured funding, project administration, manuscript revision

LF: secured funding, project administration, manuscript revision

## Notes

### Competing Interest Statement

The authors have declared no competing interest.

