## Supplementary information for "Hydrological Connectivity Enhances Fish Biodiversity in Amazonian Mining Ponds: Insights from eDNA and Traditional Sampling"

**Running title: Fish Biodiversity in Amazon Mining Ponds**

Camila Timana-Mendoza* ^1^, Alonso Reyes-Calderón ^2^, Patrick Venail ^3, 4^, Ricardo Britzke ^5^, Monica C. Santa-Maria ^2, 6^, Julio M. Araújo-Flores ^1, 7, 8^, Miles Silman^1, 8^, Luis E. Fernandez^1, 8^

^1^ Centro de Innovación Científica Amazónica - CINCIA, Puerto Maldonado, Madre de Dios 17001, Peru.

^2^ Centro de Investigación y Tecnología del Agua - CITA, Universidad de Ingenieria y Tecnologia - UTEC, Lima 15063, Peru.

^3^ Microbiology Research Center - CIMIC, Department of Biological Sciences, Universidad de los Andes, Bogotá, Colombia

^4^ Inka Terra Asociación – ITA, Calle Víctor Larco Herrera 130 Miraflores, Lima, 15074, Peru.

^5^ Departamento de Ictiología, Museo de Historia Natural, Universidad Nacional Mayor de San Marcos, Lima, Peru.

^6^ Departamento de Ingeniería Ambiental, Universidad de Ingenieria y Tecnologia - UTEC, Lima 15063, Peru.

^7^ Colección Científica de Ictiología, Universidad Nacional Amazónica de Madre de Dios, Puerto Maldonado 1160, Madre de Dios, Peru.

^8^ Sabin Center for Environment and Sustainability, Wake Forest University, 1834 Wake Forest Rd, Winston-Salem, NC 27109, USA

* Corresponding autor:

**Supplementary information 1.** Study sites physicochemical properties and volume of water filtered at each site. NFP: Non-flooded [abandoned mining] ponds. FP: Flooded [abandoned mining] ponds. OL: [pristine] Oxbow lakes.

| **Site name** | **Code** | **pH** | **Conductivity (µS/s)** | **Temperature (°C)** | **Dissolved oxygen (ppm)** | **Transparency (m)** | **Volume filtered (ml)** | |
| --- | --- | --- | --- | --- | --- | --- | --- | --- |
|  |  |  |  |  |  |  | **Rep 1** | **Rep 2** |
| Lechuza | NFP#1 | 6.93 | 2.3 | 26.6 | 6.2 | 1.627 | 950 | 950 |
| Mega | NFP#2 | 7.1 | 9.6 | 27.64 | 5.9 | 4.635 | 750 | 700 |
| Cobra | NFP#3 | 6.6 | 9 | 27.5 | 5 | 1.24 | 700 | 600 |
| Balata | NFP#4 | 6.35 | 13 | 27.3 | 5.4 | 0.645 | 600 | 750 |
| Chambira | FP#1 | 6.57 | 81.7 | 29.5 | 4.4 | 0.16 | 850 | 850 |
| Ronsoco | FP#2 | 8.3 | 10 | 24 | 6 | 0.52 | 700 | 700 |
| Nueva Charapa | FP#3 | 7.57 | 86 | 28.2 | 4.6 | 0.3125 | 400 | 400 |
| Inundación | FP#4 | 7.8 | 109.5 | 28.85 | 4 | 0.5475 | 800 | 800 |
| Shansho | FP#5 | 7.58 | 75.75 | 28.125 | 5 | 0.4 | 700 | 700 |
| Katycocha | OL#1 | 6.93 | 22 | 30.7 | 5.6 | 0.81 | 600 | 600 |
| Jimena | OL#2 | 6.93 | 52 | 30.4 | 5.6 | 0.49 | 550 | 500 |

**Supplementary information 2.** Number of reads assigned to the Actinopterygii class using the 12S marker, reported for each site, field station and laboratory controls. NFP: Non-flooded [abandoned mining] ponds. FP: Flooded [abandoned mining] ponds. OL: [pristine] Oxbow lakes.

| **Order** | **Family** | **Genus** | **Species** | **Morphospecies** | **Similarity** | **Lechuza**  **NFP#1** | **Mega**  **NFP#2** | **Cobra**  **NFP#3** | **Balata**  **NFP#4** | **Chambira**  **FP#1** | **Ronsoco**  **FP#2** | **Nueva Charapa**  **FP#3** | **Inundacion**  **FP#4** | **Shansho**  **FP#5** | **Katycocha**  **OL#1** | **Jimena**  **OL#2** | **Field station control** | **Lab control** |
| --- | --- | --- | --- | --- | --- | --- | --- | --- | --- | --- | --- | --- | --- | --- | --- | --- | --- | --- |
| Characiformes | Acestrorhynchidae | *Acestrorhynchus* | *Acestrorhynchus afflacustris* | *Acestrorhynchus aff lacustris* | 99.06 | - | 270 | - | - | - | 850 | 613 | 2105 | 4009 | 9137 | 6100 | - | - |
| Characiformes | Anostomidae | *Leporinus* |  | *Leporinus sp* |  | - | - | - | - | - | 71 | 155 | 4604 | 262 | - | 52865 | - | - |
| Characiformes | Anostomidae | *Leporinus* |  | *Leporinus sp* |  | 7054 | - | 6481 | 7114 | - | 20214 | 232 | 7210 | 2993 | 11143 | 3407 | - | - |
| Characiformes | Anostomidae | *Schizodon* |  | *Schizodon sp* |  | - | - | - | - | - | 93 | 446 | 7211 | 7300 | 6699 | 706 | - | - |
| Characiformes | Anostomidae |  |  |  |  | - | - | - | - | - | - | - | 1252 | 362 | - | - | - | - |
| Characiformes | Characidae | *Astyanax* | *Astyanax bimaculatus* | *Astyanax bimaculatus* | 99.07 | - | - | - | - | 1467 | 1940 | - | 1384 | 461 | 334 | 149 | - | - |
| Characiformes | Characidae | *Compsura* |  | *Compsura sp* |  | - | - | - | - | - | - | - | - | 143 | - | 76 | - | - |
| Characiformes | Characidae | *Ctenobrycon* | *Ctenobrycon hauxwellianus* | *Ctenobrycon hauxwellianus* | 100 | - | - | 6745 | 14764 | 4052 | 2366 | 923 | 1050 | 4213 | 334 | 91 | - | - |
| Characiformes | Characidae | *Cynopotamus* |  | *Cynopotamus sp* |  | - | - | - | - | - | - | - | 1165 | 355 | 1720 | - | - | - |
| Characiformes | Characidae | *Gymnocorymbus* | *Gymnocorymbus ternetzi* | *Gymnocorymbus ternetzi* | 99.07 | 2254 | - | 6622 | 13592 | - | 1258 | 133 | 166 | 42 | 547 | - | - | - |
| Characiformes | Characidae | *Hemigrammus* |  | *Hemigrammus sp* |  | - | - | - | - | - | 3500 | - | - | - | 462 | - | - | - |
| Characiformes | Characidae | *Moenkhausia* | *Moenkhausia sanctaefilomenae* | *Moenkhausia sanctaefilomenae* | 99.06 | - | - | - | - | - | - | - | - | - | 765 | 224 | - | - |
| Characiformes | Characidae | *Moenkhausia* |  | *Moenkhausia sp* |  | - | - | - | - | - | 406 | - | 33 | - | - | - | - | - |
| Characiformes | Characidae | *Phenacogaster* |  | *Phenacogaster sp* |  | - | - | - | - | - | - | 4071 | 633 | 1334 | - | 97 | - | - |
| Characiformes | Characidae | *Poptella* | *Poptella compressa* | *Poptella compressa* | 100 | - | - | - | - | - | - | - | - | - | 497 | - | - | - |
| Characiformes | Characidae | *Roeboides* |  | *Roeboides sp* |  | - | - | - | - | - | 624 | - | 877 | 411 | 1508 | 558 | - | - |
| Characiformes | Characidae | *Serrapinnus* | *Serrapinnus piaba* | *Serrapinnus piaba* | 100 | 48610 | - | 47977 | 66510 | 19723 | 14154 | - | - | 100 | 1899 | - | - | - |
| Characiformes | Characidae |  |  |  |  | - | - | - | - | 56662 | 97 | 417 | 3413 | 4159 | 4160 | 59 | - | - |
| Characiformes | Characidae |  |  |  |  | - | - | - | - | - | - | - | 84 | - | - | - | - | - |
| Characiformes | Characidae |  |  |  |  | - | - | - | - | - | 108 | - | - | - | - | - | - | - |
| Characiformes | Characidae |  |  |  |  | - | - | - | - | - | - | - | 338 | - | - | 100 | - | - |
| Characiformes | Characidae |  |  |  |  | 4617 | - | 777 | - | - | 104 | - | - | - | - | - | - | - |
| Characiformes | Characidae |  |  |  |  | - | - | - | - | - | - | - | 5952 | 997 | 1211 | 185 | - | - |
| Characiformes | Characidae |  |  |  |  | - | - | - | - | - | - | - | 4688 | 138 | - | - | - | - |
| Characiformes | Curimatidae | *Cyphocharax* |  | *Cyphocharax sp* |  | - | - | - | - | - | 3707 | 77 | 718 | 343 | 988 | 374 | - | - |
| Characiformes | Curimatidae |  |  |  |  | - | - | - | - | 46 | 80 | - | 32 | 137 | 326 | 480 | - | - |
| Characiformes | Cynodontidae | *Hydrolycus* | *Hydrolycus scomberoides* | *Hydrolycus scomberoides* | 99.06 | - | - | - | - | - | - | - | 764 | - | - | - | - | - |
| Characiformes | Erythrinidae | *Hoplias* | *Hoplias malabaricus* | *Hoplias malabaricus* | 100 | 2156 | - | - | 2287 | 6486 | 6285 | 836 | 2709 | 2681 | 2289 | 20497 | - | - |
| Characiformes | Iguanodectidae | *Bryconops* |  | *Bryconops sp* |  | - | - | 674 | - | - | - | - | - | - | - | - | - | - |
| Characiformes | Iguanodectidae |  |  |  |  | - | - | 164 | 1782 | - | - | - | - | - | - | - | - | - |
| Characiformes | Parodontidae |  |  |  |  | - | - | - | - | 293 | 139 | 184 | 140 | 134 | 471 | 1000 | - | - |
| Characiformes | Prochilodontidae | *Prochilodus* | *Prochilodus lineatus* | *Prochilodus lineatus* | 100 | - | 24 | - | 38 | 1802 | - | 13177 | 43327 | 15159 | 658 | 8465 | - | - |
| Characiformes | Serrasalmidae | *Serrasalmus* |  | *Serrasalmus sp* |  | - | - | - | - | - | 3287 | 1567 | 408 | 462 | 23396 | 2750 | - | - |
| Characiformes |  |  |  |  |  | - | - | - | - | 212 | 142 | 229 | 160 | 137 | 514 | 1167 | - | - |
| Characiformes |  |  |  |  |  | 52 | - | 3961 | 860 | 21285 | 20919 | 149150 | 30727 | 36296 | 105541 | 50863 | - | 22 |
| Gymnotiformes | Gymnotidae | *Gymnotus* | *Gymnotus carapo* | *Gymnotus carapo* | 99.06 | - | - | - | - | - | 90 | 59 | 2310 | 539 | 184 | 108 | - | - |
| Gymnotiformes | Hypopomidae | *Brachyhypopomus* |  | *Brachyhypopomus sp* |  | - | - | - | - | - | 19 | - | - | - | - | 81 | - | - |
| Gymnotiformes | Hypopomidae | *Brachyhypopomus* |  | *Brachyhypopomus sp* |  | - | - | - | - | - | - | - | - | - | 150 | - | - | - |
| Gymnotiformes | Sternopygidae | *Eigenmannia* |  | *Eigenmannia sp* |  | - | - | - | - | - | 592 | 404 | - | 60 | - | - | - | - |
| Gymnotiformes | Sternopygidae | *Sternopygus* | *Sternopygus macrurus* | *Sternopygus macrurus* | 100 | - | - | 1816 | 11625 | - | 342 | 178 | 3587 | 4719 | 2590 | 105 | - | - |
| Gymnotiformes | Sternopygidae |  |  |  |  | - | - | - | - | - | - | 38 | 4137 | 619 | - | 1667 | - | - |
| Gymnotiformes | Sternopygidae |  |  |  |  | - | - | - | - | - | - | - | 569 | 110 | - | - | - | - |
| Perciformes | Carangidae |  |  |  |  | - | - | - | - | 1210 | - | - | - | - | - | - | - | - |
| Perciformes | Cichlidae | *Cichlasoma* |  | *Cichlasoma sp* |  | - | - | - | - | 2726 | 65 | - | 269 | 226 | 125 | 48 | - | - |
| Perciformes | Cichlidae | *Saxatilia* | *Saxatilia lepidota* | *Saxatilia lepidota* | 99.06 | 56509 | 316 | 75541 | 116859 | 3592 | 4006 | - | 9120 | 7318 | 2011 | 192 | - | - |
| Perciformes | Cichlidae |  |  |  |  | - | - | - | - | - | - | - | - | - | - | 30 | - | - |
| Perciformes | Cichlidae |  |  |  |  | 1096 | - | 92 | - | - | 111 | - | - | - | - | - | - | - |
| Perciformes | Cichlidae |  |  |  |  | - | - | - | - | - | 1203 | - | 217 | 384 | 2463 | 832 | - | - |
| Siluriformes | Auchenipteridae | *Auchenipterus* |  | *Auchenipterus sp* |  | - | - | - | - | - | - | - | 4593 | - | - | 96 | - | - |
| Siluriformes | Auchenipteridae | *Trachelyopterus* | *Trachelyopterus galeatus* | *Trachelyopterus galeatus* | 100 | - | 273 | - | - | - | 158 | 19 | - | - | - | - | - | - |
| Siluriformes | Heptapteridae | *Pimelodella* |  | *Pimelodella sp* |  | - | - | - | - | - | - | - | 1025 | - | - | - | - | - |
| Siluriformes | Loricariidae | *Ancistrus* |  | *Ancistrus sp* |  | - | - | - | - | - | 1437 | - | 769 | 1245 | 1580 | 243 | - | - |
| Siluriformes | Loricariidae | *Hypoptopoma* |  | *Hypoptopoma sp* |  | - | - | - | - | - | 4246 | - | 1546 | 430 | 405 | 140 | - | - |
| Siluriformes | Loricariidae | *Hypostomus* |  | *Hypostomus sp* |  | - | 129 | - | - | 28214 | 2598 | 835 | 4228 | 5049 | 6107 | 58593 | - | 20 |
| Siluriformes | Loricariidae | *Loricariichthys* | *Loricariichthys platymetopon* | *Loricariichthys platymetopon* | 100 | - | - | - | - | 3520 | 2980 | 192 | 4565 | 3230 | 7145 | 1607 | - | - |
| Siluriformes | Loricariidae |  |  |  |  | - | - | - | - | - | - | 134 | - | 1034 | - | 44 | - | - |
| Siluriformes | Pimelodidae | *Hemisorubim* | *Hemisorubim platyrhynchos* | *Hemisorubim platyrhynchos* | 100 | - | - | - | - | - | - | - | 674 | - | - | - | - | - |
| Siluriformes | Pimelodidae | *Hypophthalmus* | *Hypophthalmus edentatus* | *Hypophthalmus edentatus* | 100 | - | - | - | - | - | 215 | - | - | - | 1169 | - | - | - |
| Siluriformes | Pimelodidae | *Pimelodus* |  | *Pimelodus sp* |  | - | - | - | - | - | - | 19 | 3254 | 428 | - | 42 | - | - |
| Siluriformes | Pimelodidae | *Pseudoplatystoma* | *Pseudoplatystoma tigrinum* | *Pseudoplatystoma tigrinum* | 100 | - | - | - | - | - | - | 102 | 305 | 37 | - | - | - | - |
| Siluriformes | Pimelodidae | *Pseudoplatystoma* |  | *Pseudoplatystoma sp* |  | - | - | - | - | - | - | 24 | - | - | - | - | - | - |
| Siluriformes | Pimelodidae | *Sorubim* | *Sorubim lima* | *Sorubim lima* | 100 | - | - | - | - | - | - | 28 | 932 | - | - | - | - | - |
| Synbranchiformes | Synbranchidae | *Synbranchus* | *Synbranchus marmoratus* | *Synbranchus marmoratus* | 100 | - | - | - | - | - | - | - | - | - | - | 29 | - | - |

**Supplementary information 3.** Number of reads assigned to the Actinopterygii class using the COI marker, reported for each site, field station and laboratory controls. NFP: Non-flooded [abandoned mining] ponds. FP: Flooded [abandoned mining] ponds. OL: [pristine] Oxbow lakes

| **Order** | **Family** | **Genus** | **Species** | **Morphospecie** | **Similarity** | **Lechuza**  **NFP#1** | **Mega**  **NFP#2** | **Cobra**  **NFP#3** | **Balata**  **NFP#4** | **Chambira**  **FP#1** | **Ronsoco**  **FP#2** | **Nueva Charapa**  **FP#3** | **Inundacion**  **FP#4** | **Shansho**  **FP#5** | **Katycocha**  **OL#1** | **Jimena**  **OL#2** | **Field station control** | **Lab control** |
| --- | --- | --- | --- | --- | --- | --- | --- | --- | --- | --- | --- | --- | --- | --- | --- | --- | --- | --- |
| Characiformes | Anostomidae | Leporinus | *Leporinus aff. friderici* | *Leporinus aff. friderici* | 100 | - | - | 29 | - | - | 563 | - | 69 | - | 1,072 | 535 | - | - |
| Characiformes | Anostomidae | Leporinus | *Megaleporinus trifasciatus* | *Megaleporinus trifasciatus* | 100 | - | - | - | - | - | - | 69 | 21 | - | - | 19,121 | - | - |
| Characiformes | Anostomidae | Leporinus |  | *Leporinus sp* |  | - | - | - | - | - | - | - | 71 | 64 | - | - | - | - |
| Characiformes | Anostomidae | Schizodon | *Schizodon fasciatus* | *Schizodon fasciatus* | 100 | - | - | - | - | - | 67 | 221 | 555 | 386 | 1,059 | 277 | - | - |
| Characiformes | Characidae | Aphyocharax |  | *Aphyocharax sp* |  | - | - | - | - | - | - | 40 | - | 37 | - | - | - | - |
| Characiformes | Characidae | Aphyocheirodon |  | *Aphyocheirodon sp* |  | 889 | - | 1527 | 1,722 | - | 611 | 78 | - | - | 288 | - | - | - |
| Characiformes | Characidae | Astyanax | *Astyanax bimaculatus* | *Astyanax bimaculatus* | 100 | - | - | - | - | - | 174 | - | 37 | 82 | 69 | 86 | - | - |
| Characiformes | Characidae | Astyanax/Psellogrammus | *Astyanax bimaculatus* | *Astyanax bimaculatus* | 100 | 133 | - | 591 | - | 30 | 138 | 285 | 33 | 149 | 63 | 23 | - | - |
| Characiformes | Characidae | Charax | *Charax gibbosus* | *Charax gibbosus* | 100 | - | - | - | - | - | - | 1,448 | - | - | - | 47 | - | - |
| Characiformes | Characidae | Moenkhausia |  | *Moenkhausia sp* |  | - | - | - | 51 | - | 511 | - | 115 | 195 | 682 | 538 | - | - |
| Characiformes | Characidae | Roeboides | *Roeboides descalvadensis* | *Roeboides descalvadensis* | 100 | - | - | - | - | - | - | 47 | 75 | 48 | - | - | - | - |
| Characiformes | Characidae | Roeboides |  | *Roeboides sp* |  | - | - | - | - | - | 14 | - | 63 | - | - | - | - | - |
| Characiformes | Characidae | Tetragonopterus | *Tetragonopterus argenteus* | *Tetragonopterus argenteus* | 100 | - | - | - | - | - | - | - | 33 | 43 | - | - | - | - |
| Characiformes | Characidae |  |  |  |  | - | - | - | - | - | - | 92 | - | - | - | - | - | - |
| Characiformes | Characidae |  |  |  |  | - | - | - | - | - | - | - | - | - | 410 | 502 | - | - |
| Characiformes | Characidae |  |  |  |  | - | - | - | - | - | 2,579 | - | - | - | 81 | - | - | - |
| Characiformes | Characidae |  |  |  |  | - | - | - | - | - | - | - | - | - | 366 | - | - | - |
| Characiformes | Characidae |  |  |  |  | - | - | 431 | 1,315 | - | 31 | 86 | - | 49 | - | - | - | - |
| Characiformes | Characidae |  |  |  |  | - | - | - | - | - | - | - | - | 35 | - | - | - | - |
| Characiformes | Curimatidae | Curimatella |  | *Curimatella sp* |  | - | - | - | - | - | - | 78 | - | 38 | - | - | - | - |
| Characiformes | Curimatidae | Cyphocharax |  | *Cyphocharax sp* |  | - | - | - | - | - | 344 | - | 158 | 506 | 2,828 | 1,009 | - | - |
| Characiformes | Curimatidae | Potamorhina | *Potamorhina altamazonica* | *Potamorhina altamazonica* | 100 | - | 176 | - | - | 118 | 485 | 23,600 | 216 | 124 | 1,654 | 7,769 | - | - |
| Characiformes | Curimatidae | Psectrogaster | *Psectrogaster rutiloides* | *Psectrogaster rutiloides* | 100 | - | - | - | - | - | - | 88 | - | - | - | 2,571 | - | - |
| Characiformes | Curimatidae | Steindachnerina | *Steindachnerina guentheri* | *Steindachnerina guentheri* | 100 | - | - | - | - | - | 36 | - | - | - | - | 87 | - | - |
| Characiformes | Cynodontidae | Cynodon |  | *Cynodon sp* |  | - | - | - | - | - | - | - | - | - | - | 95 | - | - |
| Characiformes | Erythrinidae | Hoplerythrinus | *Hoplerythrinus unitaeniatus* | *Hoplerythrinus unitaeniatus* | 99 | - | - | 22 | - | - | 52 | - | - | - | - | - | - | - |
| Characiformes | Erythrinidae | Hoplias | *Hoplias malabaricus* | *Hoplias malabaricus* | 100 | 108 | - | - | - | 169 | 607 | 188 | 56 | 364 | 178 | 8,174 | - | - |
| Characiformes | Prochilodontidae | Prochilodus | *Prochilodus nigricans* | *Prochilodus nigricans* | 100 | - | - | - | - | - | - | 4,962 | 679 | 686 | 66 | 4,779 | - | - |
| Characiformes | Serrasalmidae | Mylossoma | *Mylossoma albiscopum* | *Mylossoma albiscopum* | 100 | - | - | - | - | - | - | - | 131 | - | - | - | - | - |
| Characiformes | Serrasalmidae | Serrasalmus | *Serrasalmus rhombeus* | *Serrasalmus rhombeus* | 100 | - | - | - | - | - | 79 | - | - | - | 706 | 127 | - | - |
| Characiformes | Serrasalmidae |  |  |  |  | - | - | - | - | - | 634 | 775 | 18 | - | 1,749 | 2,737 | - | - |
| Characiformes | Triportheidae | Triportheus |  | *Triportheus sp* |  | - | - | - | - | - | 708 | 588 | 101 | 1,507 | 340 | 854 | - | - |
| Characiformes |  |  |  |  |  | - | - | - | - | - | - | - | - | - | 289 | - | - | - |
| Characiformes |  |  |  |  |  | - | - | - | - | - | - | - | 44 | - | - | - | - | - |
| Characiformes |  |  |  |  |  | - | - | - | - | - | 29 | 135 | - | 91 | 78 | 1,193 | - | - |
| Gymnotiformes | Gymnotidae | Gymnotus | *Gymnotus carapo* | *Gymnotus carapo* | 100 | - | - | - | - | - | 127 | 45 | - | 96 | - | 395 | - | - |
| Gymnotiformes | Hypopomidae | Brachyhypopomus | *Brachyhypopomus pinnicaudatus* | *Brachyhypopomus pinnicaudatus* | 100 | - | - | - | - | - | 73 | - | - | - | - | - | - | - |
| Gymnotiformes | Hypopomidae |  |  |  |  | - | - | - | - | - | - | - | - | - | 40 | - | - | - |
| Gymnotiformes | Sternopygidae | Eigenmannia | *Eigenmannia limbata* | *Eigenmannia limbata* | 100 | - | - | - | - | - | - | - | - | - | - | 51 | - | - |
| Gymnotiformes | Sternopygidae | Eigenmannia | *Eigenmannia gr. trilineata* | *Eigenmannia gr. trilineata* | 99 | - | - | - | - | - | - | - | - | - | - | 282 | - | - |
| Gymnotiformes | Sternopygidae | Eigenmannia |  | *Eigenmannia sp* |  | - | - | - | - | - | 151 | 262 | - | - | - | - | - | - |
| Gymnotiformes | Sternopygidae | Sternopygus | *Sternopygus macrurus* | *Sternopygus macrurus* | 100 | - | - | 548 | 428 | - | - | 100 | 33 | 308 | - | 308 | - | - |
| Perciformes | Cichlidae | Cichlasoma | *Cichlasoma bimaculatum* | *Cichlasoma bimaculatum* | 99 | - | - | - | 126 | 648 | 276 | 545 | 286 | 1,674 | 331 | 98 | - | - |
| Perciformes | Cichlidae | Cichlasoma | *Cichlasoma bimaculatum* | *Cichlasoma bimaculatum* | 99 | 5,968 | - | 547 | - | - | 512 | - | - | - | - | - | - | - |
| Perciformes | Cichlidae | Cichlasoma |  | *Cichlasoma sp* |  | - | - | - | 886 | - | - | - | - | - | - | - | - | - |
| Perciformes | Cichlidae | Crenicichla |  | *Crenicichla sp* |  | 2,512 | 1,720 | 10894 | 15,914 | 109 | 674 | - | 111 | 1,524 | - | 735 | - | - |
| Perciformes | Cichlidae | Mesonauta | *Mesonauta festivus* | *Mesonauta festivus* | 100 | - | - | - | - | - | - | - | - | 28 | 217 | - | - | - |
| Perciformes | Cichlidae | Satanoperca |  | *Satanoperca sp* |  | - | - | - | - | - | 3,998 | - | 789 | 2,766 | 4,802 | 9,519 | - | - |
| Perciformes | Cichlidae |  |  |  |  | - | - | - | - | - | 380 | 84 | 34 | 680 | - | - | - | - |
| Perciformes | Cichlidae |  |  |  |  | - | - | - | - | - | - | - | - | - | 400 | 539 | - | - |
| Perciformes | Cichlidae |  |  |  |  | - | - | - | - | - | - | - | - | 28 | - | - | - | - |
| Perciformes | Cichlidae |  |  |  |  | - | - | - | - | - | 154 | - | - | - | - | - | - | - |
| Siluriformes | Auchenipteridae | Auchenipterus | *Auchenipterus ambyiacus/Auchenipterus nuchalis* | *Auchenipterus ambyiacus/Auchenipterus nuchalis* | 100 | - | - | - | - | - | - | - | 37 | - | 262 | - | - | - |
| Siluriformes | Auchenipteridae | Trachelyopterus | *Trachelyopterus galeatus* | *Trachelyopterus galeatus* | 100 | - | - | - | - | - | 70 | - | - | - | - | - | - | - |
| Siluriformes | Loricariidae | Ancistrus |  | *Ancistrus sp* |  | - | - | - | - | - | 183 | - | - | 105 | 2,080 | 149 | - | - |
| Siluriformes | Loricariidae | Hypoptopoma | *Hypoptopoma gulare* | *Hypoptopoma gulare* | 100 | - | - | - | - | - | 499 | 74 | 24 | - | 161 | - | - | - |
| Siluriformes | Loricariidae | Hypostomus |  | *Hypostomus sp* |  | - | - | - | - | 64 | 65 | 695 | 31 | 435 | 972 | 16,376 | - | - |
| Siluriformes | Loricariidae | Loricariichthys | *Loricariichthys platymetopon* | *Loricariichthys platymetopon* | 100 | - | - | - | - | 22 | 908 | 128 | 119 | 771 | 1,402 | 1,756 | - | - |
| Siluriformes | Pimelodidae | Hypophthalmus |  | *Hypophthalmus sp* |  | - | - | - | - | - | 40 | - | - | - | 125 | - | - | - |
| Siluriformes | Pimelodidae | Pimelodus | *Pimelodus tetramerus* | *Pimelodus tetramerus* | 100 | - | - | - | - | - | - | - | 72 | 50 | - | - | - | - |
| Siluriformes | Pimelodidae | Sorubim | *Sorubim lima* | *Sorubim lima* | 100 | - | - | - | - | - | - | 50 | - | - | - | - | - | - |

**Supplementary information 4.** Family presence across study sites identified using the 12S and COI markers. '1' indicates presence, and '0' indicates absence. NFP: Non-flooded [abandoned mining] ponds. FP: Flooded [abandoned mining] ponds. OL: [pristine] Oxbow lakes

| **Family** | **Lechuza (NFP#1)**  **12S/COI** | **Mega (NFP#2)**  **12S/COI** | **Cobra (NFP#3)**  **12S/COI** | **Balata (NFP#4)**  **12S/COI** | **Chambira (FP#1)**  **12S/COI** | **Ronsoco (FP#2)**  **12S/COI** | **Ncharapa (FP#3)**  **12S/COI** | **Inundacion (FP#4)**  **12S/COI** | **Shansho (FP#5)**  **12S/COI** | **Katycocha (OL#1)**  **12S/COI** | **Jimena (OL#2)**  **12S/COI** |
| --- | --- | --- | --- | --- | --- | --- | --- | --- | --- | --- | --- |
| Acestrorhynchidae | 0/0 | 1/0 | 0/0 | 0/0 | 0/0 | 1/0 | 1/0 | 1/0 | 1/0 | 1/0 | 1/0 |
| Anostomidae | 1/0 | 0/0 | 1/1 | 1/0 | 0/0 | 1/1 | 1/1 | 1/1 | 1/1 | 1/1 | 1/1 |
| Auchenipteridae | 0/0 | 1/0 | 0/0 | 0/0 | 0/0 | 1/1 | 1/0 | 1/1 | 0/0 | 0/1 | 1/0 |
| Characidae | 1/1 | 0/0 | 1/1 | 1/1 | 1/1 | 1/1 | 1/1 | 1/1 | 1/1 | 1/1 | 1/1 |
| Characiformes* | 1/0 | 0/0 | 1/0 | 1/0 | 1/0 | 1/1 | 1/1 | 1/1 | 1/1 | 1/1 | 1/1 |
| Cichlidae | 1/1 | 1/1 | 1/1 | 1/1 | 1/1 | 1/1 | 0/1 | 1/1 | 1/1 | 1/1 | 1/1 |
| Curimatidae | 0/0 | 0/1 | 0/0 | 0/0 | 1/1 | 1/1 | 1/1 | 1/1 | 1/1 | 1/1 | 1/1 |
| Cynodontidae | 0/0 | 0/0 | 0/0 | 0/0 | 0/0 | 0/0 | 0/0 | 1/0 | 0/0 | 0/0 | 0/1 |
| Erythrinidae | 1/1 | 0/0 | 0/1 | 1/0 | 1/1 | 1/1 | 1/1 | 1/1 | 1/1 | 1/1 | 1/1 |
| Gymnotidae | 0/0 | 0/0 | 0/0 | 0/0 | 0/0 | 1/1 | 1/1 | 1/0 | 1/1 | 1/0 | 1/1 |
| Heptapteridae | 0/0 | 0/0 | 0/0 | 0/0 | 0/0 | 0/0 | 0/0 | 1/0 | 0/0 | 0/0 | 0/0 |
| Hypopomidae | 0/0 | 0/0 | 0/0 | 0/0 | 0/0 | 1/1 | 0/0 | 0/0 | 0/0 | 1/1 | 1/0 |
| Iguanodectidae | 0/0 | 0/0 | 1/0 | 1/0 | 0/0 | 0/0 | 0/0 | 0/0 | 0/0 | 0/0 | 0/0 |
| Loricariidae | 0/0 | 1/0 | 0/0 | 0/0 | 1/1 | 1/1 | 1/1 | 1/1 | 1/1 | 1/1 | 1/1 |
| Parodontidae | 0/0 | 0/0 | 0/0 | 0/0 | 1/0 | 1/0 | 1/0 | 1/0 | 1/0 | 1/0 | 1/0 |
| Pimelodidae | 0/0 | 0/0 | 0/0 | 0/0 | 0/0 | 1/1 | 1/1 | 1/1 | 1/1 | 1/1 | 1/0 |
| Prochilodontidae | 0/0 | 1/0 | 0/0 | 1/0 | 1/0 | 0/0 | 1/1 | 1/1 | 1/1 | 1/1 | 1/1 |
| Serrasalmidae | 0/0 | 0/0 | 0/0 | 0/0 | 0/0 | 1/1 | 1/1 | 1/1 | 1/0 | 1/1 | 1/1 |
| Sternopygidae | 0/0 | 0/0 | 1/1 | 1/1 | 0/0 | 1/1 | 1/1 | 1/1 | 1/1 | 1/0 | 1/1 |
| Synbranchidae | 0/0 | 0/0 | 0/0 | 0/0 | 0/0 | 0/0 | 0/0 | 0/0 | 0/0 | 0/0 | 1/0 |
| Triportheidae | 0/0 | 0/0 | 0/0 | 0/0 | 0/0 | 0/1 | 0/1 | 0/1 | 0/1 | 0/1 | 0/1 |

**Supplementary information 5.** Taxa recorded using traditional methods (TM). Numbers in each cell represent the number of individuals per species. NFP: Non-flooded [abandoned mining] ponds. FP: Flooded [abandoned mining] ponds. OL: [pristine] Oxbow lakes

| **Order** | **Family** | **Genus** | **Species** | **Morphospecie** | **Lechuza**  **NFP#1** | **Mega**  **NFP#2** | **Cobra**  **NFP#3** | **Balata**  **NFP#4** | **Chambira**  **FP#1** | **Ronsoco**  **FP#2** | **Nueva Charapa**  **FP#3** | **Inundacion**  **FP#4** | **Shansho**  **FP#5** | **Katycocha**  **OL#1** | **Jimena**  **OL#2** |
| --- | --- | --- | --- | --- | --- | --- | --- | --- | --- | --- | --- | --- | --- | --- | --- |
| Characiformes | Hemiodontidae | Anodus | *Anodus elongatus* | *Anodus elongatus* | - | - | - | - | - | - | - | 3 | - | - | - |
| Characiformes | Curimatidae | Cyphocharax | *Cyphocharax spiluropsis* | *Cyphocharax spiluropsis* | - | - | - | - | - | 1 | - | 18 | - | 8 | - |
| Characiformes | Curimatidae | Curimatella | *Curimatella meyeri* | *Curimatella meyeri* | - | - | - | - | - | - | - | 1 | - | - | - |
| Characiformes | Curimatidae | Steindachnerina | *Steindachnerina guentheri* | *Steindachnerina guentheri* | - | - | - | - | 30 | 5 | - | - | - | 1 | - |
| Characiformes | Curimatidae | Steindachnerina | *Steindachnerina aff. dobula* | *Steindachnerina aff. dobula* | - | - | - | - | 23 | 5 | - | 18 | 5 | - | - |
| Characiformes | Curimatidae | Steindachnerina |  | *Steindachnerina sp* | - | - | - | - | - | - | - | 1 | - | - | - |
| Characiformes | Curimatidae | Potamorhina | *Potamorhina altamazonica* | *Potamorhina altamazonica* | - | - | - | - | - | 4 | 5 | 2 | 4 | 16 | - |
| Characiformes | Curimatidae | Psectrogaster | *Psectrogaster rutiloides* | *Psectrogaster rutiloides* | - | - | - | - | 5 | - | - | 3 | - | - | - |
| Characiformes | Prochilodontidae | Prochilodus | *Prochilodus nigricans* | *Prochilodus nigricans* | - | - | - | - | - | 18 | 4 | 3 | - | 1 | - |
| Characiformes | Anostomidae | Leporinus | *Leporinus cf. subniger* | *Leporinus cf. subniger* | 1 | - | 3 | - | 1 | - | - | 5 | 1 | 1 | - |
| Characiformes | Anostomidae | Schizodon | *Schizodon fasciatus* | *Schizodon fasciatus* | - | - | - | 1 | - | - | - | 1 | - | - | - |
| Characiformes | Characidae | Aphyocharax | *Aphyocharax avary* | *Aphyocharax avary* | - | - | - | - | 9 | 2 | 6 | 4 | 3 | 5 | - |
| Characiformes | Characidae | Astyanax | *Astyanax bimaculatus* | *Astyanax bimaculatus* | - | - | - | - | 3 | 2 | - | 16 | 1 | 10 | - |
| Characiformes | Characidae | Charax | *Charax gibbosus* | *Charax gibbosus* | - | - | - | - | - | 1 | 25 | - | 1 | - | 6 |
| Characiformes | Characidae | Charax | *Charax pauciradiatus* | *Charax pauciradiatus* | - | - | - | - | - | - | - | 26 | - | - | - |
| Characiformes | Characidae | Ctenobrycon | *Ctenobrycon hauxwellianus* | *Ctenobrycon hauxwellianus* | - | - | - | - | 21 | 15 | 96 | 7 | 1 | 7 | 12 |
| Characiformes | Characidae | Galeocharax | *Galeocharax gulo* | *Galeocharax gulo* | - | - | - | - | - | - | - | 1 | - | - | - |
| Characiformes | Characidae | Gymnocorymbus | *Gymnocorymbus thayeri* | *Gymnocorymbus thayeri* | - | - | - | - | - | - | - | - | - | - | 7 |
| Characiformes | Characidae | Gymnocorymbus | *Gymnocorymbus ternetzi* | *Gymnocorymbus ternetzi* | - | - | - | - | - | - | - | - | - | - | 2 |
| Characiformes | Characidae | Brachychalcinus | *Brachychalcinus aff. nummus* | *Brachychalcinus aff. nummus* | - | - | - | - | - | 8 | - | - | - | - | - |
| Characiformes | Characidae |  |  |  | - | - | - | - | - | - | - | - | - | 2 | - |
| Characiformes | Characidae | Brachychalcinus | *Brachychalcinus copei* | *Brachychalcinus copei* | - | - | - | 4 | - | - | 6 | - | - | - | - |
| Characiformes | Characidae | Hemigrammus |  | *Hemigrammus sp* | - | - | - | - | - | - | - | 4 | 4 | - | 5 |
| Characiformes | Characidae | Jupiaba | *Jupiaba anteroides* | *Jupiaba anteroides* | - | - | - | - | - | - | - | - | - | - | 1 |
| Characiformes | Characidae | Knodus |  | *Knodus sp. 1* | - | - | - | - | - | - | - | - | - | - | 33 |
| Characiformes | Characidae | Knodus |  | *Knodus sp. 2* | - | - | - | - | - | - | - | - | - | - | 94 |
| Characiformes | Characidae | Knodus |  | *Knodus sp. 3* | - | - | - | - | - | - | - | - | - | - | 19 |
| Characiformes | Characidae | Knodus |  | *Knodus sp. 4* | - | - | - | - | - | - | - | - | - | - | 2 |
| Characiformes | Characidae | Knodus |  | *Knodus sp. 5* | - | 4 | - | 7 | - | - | - | - | - | 2 | - |
| Characiformes | Characidae | Knodus |  | *Knodus sp. 6* | - | - | - | - | 157 | 195 | 7 | 38 | - | - | - |
| Characiformes | Characidae | Moenkhausia | *Moenkhausia madeirae* | *Moenkhausia madeirae* | - | - | - | - | 1 | 85 | 19 | 74 | 64 | 10 | 7 |
| Characiformes | Characidae | Moenkhausia | *Moenkhausia bonita* | *Moenkhausia bonita* | - | - | - | - | - | - | - | - | - | - | 10 |
| Characiformes | Characidae | Moenkhausia | *Moenkhausia lepidura* | *Moenkhausia lepidura* | - | - | - | 1 | - | - | - | - | - | - | - |
| Characiformes | Characidae | Moenkhausia | *Moenkhausia oligolepis* | *Moenkhausia oligolepis* | - | - | - | - | - | 2 | - | - | - | - | - |
| Characiformes | Characidae | Roeboides | *Roeboides biserialis* | *Roeboides biserialis* | - | - | - | - | - | - | - | 7 | - | - | - |
| Characiformes | Characidae | Roeboides | *Roeboides afinnis* | *Roeboides afinnis* | - | - | - | - | - | - | 4 | - | - | - | - |
| Characiformes | Characidae | Serrapinnus |  | *Serrapinnus sp. 1* | - | - | - | - | 39 | 31 | 11 | 28 | 17 | 160 | - |
| Characiformes | Characidae | Serrapinnus |  | *Serrapinnus sp. 2* | - | - | - | - | - | - | 19 | - | - | - | - |
| Characiformes | Characidae | Tetragonopterus | *Tetragonopterus argenteus* | *Tetragonopterus argenteus* | - | - | - | - | - | - | - | 3 | - | - | - |
| Characiformes | Triportheidae | Triportheus | *Triportheus angulatus* | *Triportheus angulatus* | - | - | - | - | - | - | 5 | 22 | - | 5 | 3 |
| Characiformes | Triportheidae | Triportheus | *Triportheus rotundatus* | *Triportheus rotundatus* | - | - | - | - | 3 | - | - | - | - | - | - |
| Characiformes | Characidae | Tyttobrycon |  | *Tyttobrycon sp* | - | - | - | - | - | 11 | - | - | - | - | - |
| Characiformes | Characidae |  |  |  | - | - | - | - | - | - | - | - | - | - | 1 |
| Characiformes | Crenuchidae | Characidium |  | *Characidium sp* | - | - | - | - | - | - | 1 | - | - | - | - |
| Characiformes | Erythrinidae | Hoplias | *Hoplias malabaricus* | *Hoplias malabaricus* | - | 9 | 3 | 1 | 9 | 1 | 2 | 9 | 4 | 4 | - |
| Characiformes | Erythrinidae | Hoplerythrinus | *Hoplerythrinus unitaeniatus* | *Hoplerythrinus unitaeniatus* | 2 | - | 1 | - | - | - | - | - | - | - | - |
| Characiformes | Bryconidae | Bryconops | *Bryconops melanurus* | *Bryconops melanurus* | - | - | - | 2 | - | - | - | - | - | - | - |
| Characiformes | Cynodontidae | Cynodon | *Cynodon gibbus* | *Cynodon gibbus* | - | - | - | - | - | - | - | 1 | - | - | - |
| Characiformes | Acestrorhynchidae | Acestrorhynchus | *Acestrorhynchus falcatus* | *Acestrorhynchus falcatus* | - | - | - | - | - | - | - | 13 | 1 | 1 | 2 |
| Characiformes | Serrasalmidae | Mylossoma | *Mylossoma albiscopum* | *Mylossoma albiscopum* | - | - | - | - | - | - | - | 2 | - | - | - |
| Characiformes | Serrasalmidae | Serrasalmus | *Serrasalmus rhombeus* | *Serrasalmus rhombeus* | - | - | - | - | - | 5 | 1 | - | - | - | 6 |
| Characiformes | Serrasalmidae | Serrasalmus | *Serrasalmus maculatus* | *Serrasalmus maculatus* | - | - | - | - | - | 3 | 1 | 1 | - | 2 | 8 |
| Characiformes | Serrasalmidae | Serrasalmus | *Serrasalmus spilopleura* | *Serrasalmus spilopleura* | - | - | - | - | - | - | - | 1 | - | - | 5 |
| Siluriformes | Auchenipteridae | Trachelyopterus | *Trachelyopterus porosus* | *Trachelyopterus porosus* | - | - | - | 2 | - | - | - | - | - | - | - |
| Siluriformes | Auchenipteridae | Auchenipterus | *Auchenipterus ambyiacus* | *Auchenipterus ambyiacus* | - | - | - | 1 | - | - | - | 1 | - | - | - |
| Siluriformes | Loricariidae | Hypostomus |  | *Hypostomus sp.1* | - | - | - | - | - | 1 | 6 | 2 | 3 | 7 | - |
| Siluriformes | Loricariidae | Hypostomus |  | *Hypostomus sp.2* | - | - | - | - | - | - | 1 | 1 | - | - | - |
| Siluriformes | Loricariidae | Loricarichthys |  | *Loricarichthys sp* | - | - | - | - | - | - | - | - | - | 2 | - |
| Siluriformes | Loricariidae | Loricarichthys | *Loricariichthys platymetopon* | *Loricariichthys platymetopon* | - | - | - | - | 8 | 42 | - | 1 | - | - | 3 |
| Siluriformes | Loricariidae | Pterygoplichthys | *Pterygoplichthys disjunctivus* | *Pterygoplichthys disjunctivus* | - | - | - | - | - | - | 2 | - | - | - | - |
| Siluriformes | Loricariidae | Hypoptopoma | *Hypoptopoma gulare* | *Hypoptopoma gulare* | - | - | - | - | - | - | - | - | - | - | 1 |
| Siluriformes | Loricariidae | Hypoptopoma | *Hypoptopoma cf. bianale* | *Hypoptopoma cf. bianale* | - | - | - | - | - | - | - | - | 1 | - | - |
| Siluriformes | Pimelodidae | Hypophthalmus | *Hypophthalmus edentatus* | *Hypophthalmus edentatus* | - | - | - | - | - | - | - | - | - | - | 2 |
| Siluriformes | Pimelodidae | Sorubim | *Sorubim lima* | *Sorubim lima* | - | - | - | - | - | - | - | 1 | - | - | - |
| Siluriformes | Cetopsidae | Cetopsis | *Cetopsis coecutiens* | *Cetopsis coecutiens* | - | - | - | - | - | - | - | - | - | - | 1 |
| Gymnotiformes | Sternopygidae | Eigenmannia | *Eigenmannia virescens* | *Eigenmannia virescens* | - | - | - | - | - | 9 | - | - | - | - | - |
| Gymnotiformes | Sternopygidae | Eigenmannia | *Eigenmannia cf. macrops* | *Eigenmannia cf. macrops* | - | - | - | - | - | - | 2 | - | - | - | - |
| Gymnotiformes | Sternopygidae | Eigenmannia |  | *Eigenmannia sp* | - | - | - | - | - | - | - | 1 | - | - | - |
| Gymnotiformes | Sternopygidae | Sternopygus | *Sternopygus macrurus* | *Sternopygus macrurus* | - | - | - | 2 | - | - | - | 1 | - | - | - |
| Perciformes | Cichlidae | Aequidens | *Aequidens tetramerus* | *Aequidens tetramerus* | - | 44 | 1 | - | - | - | - | - | - | - | - |
| Perciformes | Cichlidae | Apistogramma |  | *Apistogramma sp* | - | - | - | - | - | - | - | 4 | 1 | - | 1 |
| Perciformes | Cichlidae | Bujurquina | *Bujurquina eurhinus* | *Bujurquina eurhinus* | - | - | - | - | 1 | 160 | - | - | - | - | - |
| Perciformes | Cichlidae | Cichlasoma | *Cichlasoma boliviense* | *Cichlasoma boliviense* | 11 | - | - | 1 | 1 | - | 1 | 3 | - | - | - |
| Perciformes | Cichlidae | Saxatilia | *Saxatilia semicincta* | *Saxatilia semicincta* | 109 | 17 | 33 | 34 | - | 1 | - | - | - | - | - |
| Perciformes | Cichlidae | Mesonauta | *Mesonauta festivus* | *Mesonauta festivus* | - | - | - | - | - | - | - | - | - | 1 | 2 |
| Perciformes | Cichlidae | Satanoperca | *Satanoperca jurupari* | *Satanoperca jurupari* | - | - | - | - | - | 8 | - | 2 | 1 | 144 | 43 |
| Beloniformes | Belonidae | Potamorrhaphis | *Potamorrhaphis cf. eigenmanni* | *Potamorrhaphis cf. eigenmanni* | - | - | - | - | - | - | - | - | - | 14 | - |
| Synbranchiformes | Synbranchidae | Synbranchus | *Synbranchus marmoratus* | *Synbranchus marmoratus* | - | - | - | - | - | - | - | - | - | 3 | - |

**Supplementary information 6.** Number of reads assigned to the Non-Actinopterygii class using the COI marker. NFP: Non-flooded [abandoned mining] ponds. FP: Flooded [abandoned mining] ponds. OL: [pristine] Oxbow lakes

| **Class** | **Order** | **Family** | **Genus** | **Species** | **Similarity** | **IUCN Red List** | **Lechuza**  **NFP#1** | **Mega**  **NFP#2** | **Cobra**  **NFP#3** | **Balata**  **NFP#4** | **Chambira**  **FP#1** | **Ronsoco**  **FP#2** | **Nueva Charapa**  **FP#3** | **Inundacion**  **FP#4** | **Shansho**  **FP#5** | **Katycocha**  **OL#1** | **Jimena**  **OL#2** |
| --- | --- | --- | --- | --- | --- | --- | --- | --- | --- | --- | --- | --- | --- | --- | --- | --- | --- |
| Amphibia | Anura | Bufonidae | Rhinella | *Rhinella marina* | 100 | Least Concern | - | - | - | - | - | - | - | - | - | - | 419 |
| Amphibia | Anura | Hylidae | Boana | *Boana geographica* | 99.23 | Least Concern | - | - | - | - | - | - | - | - | 30 | 87 | - |
| Amphibia | Anura | Hylidae | Boana |  |  |  | - | - | - | - | - | - | - | - | - | - | 47 |
| Amphibia | Anura | Hylidae | Scinax | *Scinax ruber* | 99.23 | Least Concern | - | - | - | - | - | - | - | - | 688 | - | - |
| Aves | Apodiformes | Apodidae | Chaetura | *Chaetura brachyura* | 100 | Least Concern | - | - | - | - | - | - | - | - | 36 | - | - |
| Aves | Opisthocomiformes | Opisthocomidae | Opisthocomus | *Opisthocomus hoazin* | 100 | Least Concern | - | - | - | - | - | - | - | - | 38 | - | - |
| Aves | Suliformes | Phalacrocoracidae | Phalacrocorax | *Phalacrocorax brasilianus* | 100 | Least Concern (Nannopterum brasilianus) | - | - | - | - | - | - | - | - | - | 554 | - |
| Elasmobranchii | Myliobatiformes | Potamotrygonidae | Potamotrygon |  |  |  | - | - | - | - | - | - | - | 25 | - | - | - |
| Mammalia | Perissodactyla | Tapiridae | Tapirella/Tapirus | *Tapirella bairdii/Tapirus terrestris* | 100 | Not found (*Tapirella bairdii*)/Vulnerable (*Tapirus terrestris*) | - | - | - | - | - | - | - | - | - | - | 92 |
| Reptilia | Testudines | Podocnemididae | Podocnemis | *Podocnemis unifilis* | 99.23 | Vulnerable | 685 | - | - | - | - | - | - | 33 | - | - | - |

**Supplementary information 7.** Number of reads assigned to the Non-Actinopterygii class using the 12S marker. NFP: Non-flooded [abandoned mining] ponds. FP: Flooded [abandoned mining] ponds. OL: [pristine] Oxbow lakes

| **Class** | **Order** | **Family** | **Genus** | **Species** | **Similarity** | **IUCN Red List** | **Lechuza**  **NFP#1** | **Mega**  **NFP#2** | **Cobra**  **NFP#3** | **Balata**  **NFP#4** | **Chambira**  **FP#1** | **Ronsoco**  **FP#2** | **Nueva Charapa**  **FP#3** | **Inundacion**  **FP#4** | **Shansho**  **FP#5** | **Katycocha**  **OL#1** | **Jimena**  **OL#2** |
| --- | --- | --- | --- | --- | --- | --- | --- | --- | --- | --- | --- | --- | --- | --- | --- | --- | --- |
| Amphibia | Anura | Bufonidae | Rhinella | *Rhinella marina/Rhinella poeppigii* | 100 | Least Concern (*Rhinella marina*)/Least Concern (*Rhinella poeppigii*) | - | - | - | - | - | - | - | - | - | 111 | 3999 |
| Amphibia | Anura | Bufonidae |  |  |  |  | - | - | - | - | - | - | - | - | - | 197 | - |
| Amphibia | Anura | Hylidae | Scinax |  |  |  | - | - | - | - | - | - | - | - | 40 | - | - |
| Amphibia | Anura | Leptodactylidae | Adenomera |  |  |  | - | - | - | - | - | 60 | - | - | - | - | - |
| Amphibia | Anura | Leptodactylidae | Leptodactylus |  |  |  | - | - | - | - | - | - | - | 692 | - | - | - |
| Amphibia | Anura | Phyllomedusidae | Phyllomedusa |  |  |  | - | - | - | - | - | - | - | 33 | - | - | - |
| Aves | Charadriiformes |  |  |  |  |  | - | - | - | - | - | - | - | - | - | - | 303 |
| Aves | Coraciiformes | Alcedinidae | Megaceryle |  |  |  | - | - | - | - | - | - | - | 190 | - | - | - |
| Aves | Galliformes | Cracidae |  |  |  |  | - | - | - | - | - | 228 | - | - | - | - | - |
| Aves | Opisthocomiformes | Opisthocomidae | Opisthocomus | *Opisthocomus hoazin* | 100 | Least Concern | - | - | - | - | - | - | - | - | 50 | - | - |
| Aves | Passeriformes | Hirundinidae |  |  |  |  | - | - | - | - | - | - | - | 515 | - | 1004 | - |
| Aves | Passeriformes | Thamnophilidae |  |  |  |  | - | - | - | - | - | - | - | - | 159 | - | - |
| Aves | Pelecaniformes | Ardeidae | Ardea |  |  |  | - | - | - | - | - | - | 61 | - | - | - | - |
| Aves | Psittaciformes | Psittacidae | Brotogeris |  |  |  | - | - | - | - | - | - | 24 | - | - | - | - |
| Aves |  |  |  |  |  |  | - | - | - | - | - | - | - | - | 71 | - | - |
| Mammalia | Carnivora | Mustelidae | Lontra | *Lontra longicaudis* | 100 | Near Threatened | - | - | 3090 | - | - | - | - | - | - | - | - |
| Mammalia | Chiroptera | Molossidae | Eumops |  |  |  | - | - | - | - | - | - | 19 | - | - | - | - |
| Mammalia | Chiroptera | Molossidae | Molossus | *Molossus molossus* | 100 | Least Concern | - | 99 | - | - | - | - | - | - | - | - | - |
| Mammalia | Chiroptera | Phyllostomidae | Artibeus | *Artibeus jamaicensis/Artibeus planirostris* | 100 | Least Concern (*Artibeus jamaicensis*)/Least Concern (*Artibeus planirostris*) | - | - | - | - | - | - | - | - | 60 | - | - |
| Mammalia | Chiroptera | Phyllostomidae | Artibeus |  |  |  | - | - | - | - | - | - | - | 221 | - | - | - |
| Mammalia | Perissodactyla | Tapiridae | Tapirella/Tapirus | *Tapirella bairdii/Tapirus terrestris* | 100 | Not found (*Tapirella bairdii*)/Vulnerable (*Tapirus terrestris*) | 376 | - | 1486 | - | - | 111 | - | - | - | - | 187 |
| Mammalia | Perissodactyla | Tapiridae |  |  |  |  | - | - | - | - | - | - | - | - | - | 41 | - |
| Mammalia | Primates | Atelidae | Alouatta | *Alouatta seniculus* | 100 | Least Concern | - | 251 | - | - | - | - | - | - | - | - | - |
| Mammalia | Rodentia | Caviidae | Hydrochoerus | *Hydrochoerus hydrochaeris* | 100 | Least Concern | - | - | - | - | - | 508 | - | 33 | 176 | - | 43 |
| Mammalia | Rodentia | Echimyidae | Proechimys | *Proechimys steerei* | 100 | Least Concern | - | - | - | - | - | - | - | - | - | 72 | - |
| Mammalia | Rodentia | Muridae | Rattus | *Rattus rattus* | 100 | Least Concern | - | - | - | - | - | - | - | 298 | - | - | - |
| Reptilia | Crocodylia | Alligatoridae | Melanosuchus | *Melanosuchus niger* | 100 | Lower Risk/conservation dependent | - | - | - | - | - | - | - | - | - | - | 108 |
| Reptilia | Testudines | Chelidae | Phrynops |  |  |  | - | - | - | - | - | - | - | - | 123 | - | - |
| Reptilia | Testudines | Podocnemididae | Podocnemis | *Podocnemis unifilis* | 100 | Vulnerable | - | - | - | - | 2452 | 72 | - | 99 | 276 | 810 | 407 |

**Supplementary information 8.** Taxa detected across all sites by both approaches and markers. '1' indicates presence, and '0' indicates absence. NFP: Non-flooded [abandoned mining] ponds. FP: Flooded [abandoned mining] ponds. OL: [pristine] Oxbow lakes

| **Family** | **Genus** | **Morphospecies** | **Lechuza (NFP#1)**  **12S/COI/TM** | **Mega (NFP#2)**  **12S/COI/TM** | **Cobra (NFP#3)**  **12S/COI/TM** | **Balata (NFP#4)**  **12S/COI/TM** | **Chambira (FP#1)**  **12S/COI/TM** | **Ronsoco (FP#2)**  **12S/COI/TM** | **Ncharapa (FP#3)**  **12S/COI/TM** | **Inundacion (FP#4)**  **12S/COI/TM** | **Shansho (FP#5)**  **12S/COI/TM** | **Jimena (OL#1)**  **12S/COI/TM** | **Katycocha (OL#2)**  **12S/COI/TM** |
| --- | --- | --- | --- | --- | --- | --- | --- | --- | --- | --- | --- | --- | --- |
| Acestrorhynchidae | Acestrorhynchus | *Acestrorhynchus* aff. *lacustris* | 0/0/0 | **1**/0/0 | 0/0/0 | 0/0/0 | 0/0/0 | **1**/0/0 | **1**/0/0 | **1**/0/0 | **1**/0/0 | **1**/0/0 | **1**/0/0 |
| Acestrorhynchidae | Acestrorhynchus | *Acestrorhynchus falcatus* | 0/0/0 | 0/0/0 | 0/0/0 | 0/0/0 | 0/0/0 | 0/0/0 | 0/0/0 | 0/0/**1** | 0/0/**1** | 0/0/**1** | 0/0/**1** |
| Anostomidae | Leporinus | *Leporinus* aff. *friderici* | 0/0/0 | 0/0/0 | 0/**1**/0 | 0/0/0 | 0/0/0 | 0/**1**/0 | 0/0/0 | 0/**1**/0 | 0/0/0 | 0/**1**/0 | 0/**1**/0 |
| Anostomidae | Leporinus | *Leporinus* cf. *subniger* | 0/0/**1** | 0/0/0 | 0/0/**1** | 0/0/0 | 0/0/**1** | 0/0/0 | 0/0/0 | 0/0/**1** | 0/0/**1** | 0/0/**1** | 0/0/0 |
| Anostomidae | Leporinus | *Leporinus* sp. | **1**/0/0 | 0/0/0 | **1**/0/0 | **1**/0/0 | 0/0/0 | **1**/0/0 | **1**/0/0 | **1**/**1**/0 | **1**/**1**/0 | **1**/0/0 | **1**/0/0 |
| Anostomidae | Megaleporinus | *Megaleporinus trifasciatus* | 0/0/0 | 0/0/0 | 0/0/0 | 0/0/0 | 0/0/0 | 0/0/0 | 0/**1**/0 | 0/**1**/0 | 0/0/0 | 0/**1**/0 | 0/0/0 |
| Anostomidae | Schizodon | *Schizodon fasciatus* | 0/0/0 | 0/0/0 | 0/0/0 | 0/0/**1** | 0/0/0 | 0/**1**/0 | 0/**1**/0 | 0/**1**/**1** | 0/**1**/0 | 0/**1**/0 | 0/**1**/0 |
| Anostomidae | Schizodon | *Schizodon* sp. | 0/0/0 | 0/0/0 | 0/0/0 | 0/0/0 | 0/0/0 | **1**/0/0 | **1**/0/0 | **1**/0/0 | **1**/0/0 | **1**/0/0 | **1**/0/0 |
| Auchenipteridae | Auchenipterus | *Auchenipterus ambyiacus* | 0/0/0 | 0/0/0 | 0/0/0 | 0/0/**1** | 0/0/0 | 0/0/0 | 0/0/0 | 0/0/**1** | 0/0/0 | 0/0/0 | 0/0/0 |
| Auchenipteridae | Auchenipterus | *Auchenipterus ambyiacus/Auchenipterus nuchalis* | 0/0/0 | 0/0/0 | 0/0/0 | 0/0/0 | 0/0/0 | 0/0/0 | 0/0/0 | 0/**1**/0 | 0/0/0 | 0/0/0 | 0/**1**/0 |
| Auchenipteridae | Auchenipterus | *Auchenipterus* sp. | 0/0/0 | 0/0/0 | 0/0/0 | 0/0/0 | 0/0/0 | 0/0/0 | 0/0/0 | **1**/0/0 | 0/0/0 | **1**/0/0 | 0/0/0 |
| Auchenipteridae | Trachelyopterus | *Trachelyopterus galeatus* | 0/0/0 | **1**/0/0 | 0/0/0 | 0/0/0 | 0/0/0 | **1**/**1**/0 | **1**/0/0 | 0/0/0 | 0/0/0 | 0/0/0 | 0/0/0 |
| Auchenipteridae | Trachelyopterus | *Trachelyopterus porosus* | 0/0/0 | 0/0/0 | 0/0/0 | 0/0/**1** | 0/0/0 | 0/0/0 | 0/0/0 | 0/0/0 | 0/0/0 | 0/0/0 | 0/0/0 |
| Belonidae | Potamorrhaphis | *Potamorrhaphis* cf. *eigenmanni* | 0/0/0 | 0/0/0 | 0/0/0 | 0/0/0 | 0/0/0 | 0/0/0 | 0/0/0 | 0/0/0 | 0/0/0 | 0/0/**1** | 0/0/0 |
| Cetopsidae | Cetopsis | *Cetopsis coecutiens* | 0/0/0 | 0/0/0 | 0/0/0 | 0/0/0 | 0/0/0 | 0/0/0 | 0/0/0 | 0/0/0 | 0/0/0 | 0/0/0 | 0/0/**1** |
| Characidae | Aphyocharax | *Aphyocharax avary* | 0/0/0 | 0/0/0 | 0/0/0 | 0/0/0 | 0/0/**1** | 0/0/**1** | 0/0/**1** | 0/0/**1** | 0/0/**1** | 0/0/**1** | 0/0/0 |
| Characidae | Aphyocharax | *Aphyocharax* sp. | 0/0/0 | 0/0/0 | 0/0/0 | 0/0/0 | 0/0/0 | 0/0/0 | 0/**1**/0 | 0/0/0 | 0/**1**/0 | 0/0/0 | 0/0/0 |
| Characidae | Aphyocheirodon | *Aphyocheirodon* sp. | 0/**1**/0 | 0/0/0 | 0/**1**/0 | 0/**1**/0 | 0/0/0 | 0/**1**/0 | 0/**1**/0 | 0/0/0 | 0/0/0 | 0/0/0 | 0/**1**/0 |
| Characidae | Astyanax | *Astyanax bimaculatus* | 0/**1**/0 | 0/0/0 | 0/**1**/0 | 0/0/0 | **1**/**1**/**1** | **1**/**1**/**1** | 0/**1**/0 | **1**/**1**/**1** | **1**/**1**/**1** | **1**/**1**/**1** | **1**/**1**/0 |
| Characidae | Brachychalcinus | *Brachychalcinus* aff. *nummus* | 0/0/0 | 0/0/0 | 0/0/0 | 0/0/0 | 0/0/0 | 0/0/**1** | 0/0/0 | 0/0/0 | 0/0/0 | 0/0/0 | 0/0/0 |
| Characidae | Brachychalcinus | *Brachychalcinus copei* | 0/0/0 | 0/0/0 | 0/0/0 | 0/0/**1** | 0/0/0 | 0/0/0 | 0/0/**1** | 0/0/0 | 0/0/0 | 0/0/0 | 0/0/0 |
| Characidae | Charax | *Charax gibbosus* | 0/0/0 | 0/0/0 | 0/0/0 | 0/0/0 | 0/0/0 | 0/0/**1** | 0/**1**/**1** | 0/0/0 | 0/0/**1** | 0/**1**/0 | 0/0/**1** |
| Characidae | Charax | *Charax pauciradiatus* | 0/0/0 | 0/0/0 | 0/0/0 | 0/0/0 | 0/0/0 | 0/0/0 | 0/0/0 | 0/0/**1** | 0/0/0 | 0/0/0 | 0/0/0 |
| Characidae | Compsura | *Compsura* sp. | 0/0/0 | 0/0/0 | 0/0/0 | 0/0/0 | 0/0/0 | 0/0/0 | 0/0/0 | 0/0/0 | **1**/0/0 | **1**/0/0 | 0/0/0 |
| Characidae | Ctenobrycon | *Ctenobrycon hauxwellianus* | 0/0/0 | 0/0/0 | **1**/0/0 | **1**/0/0 | **1**/0/**1** | **1**/0/**1** | **1**/0/**1** | **1**/0/**1** | **1**/0/**1** | **1**/0/**1** | **1**/0/**1** |
| Characidae | Cynopotamus | *Cynopotamus* sp. | 0/0/0 | 0/0/0 | 0/0/0 | 0/0/0 | 0/0/0 | 0/0/0 | 0/0/0 | **1**/0/0 | **1**/0/0 | 0/0/0 | **1**/0/0 |
| Characidae | Galeocharax | *Galeocharax gulo* | 0/0/0 | 0/0/0 | 0/0/0 | 0/0/0 | 0/0/0 | 0/0/0 | 0/0/0 | 0/0/**1** | 0/0/0 | 0/0/0 | 0/0/0 |
| Characidae | Gymnocorymbus | *Gymnocorymbus ternetzi* | **1**/0/0 | 0/0/0 | **1**/0/0 | **1**/0/0 | 0/0/0 | **1**/0/0 | **1**/0/0 | **1**/0/0 | **1**/0/0 | 0/0/0 | **1**/0/**1** |
| Characidae | Gymnocorymbus | *Gymnocorymbus thayeri* | 0/0/0 | 0/0/0 | 0/0/0 | 0/0/0 | 0/0/0 | 0/0/0 | 0/0/0 | 0/0/0 | 0/0/0 | 0/0/0 | 0/0/**1** |
| Characidae | Hemigrammus | *Hemigrammus* sp. | 0/0/0 | 0/0/0 | 0/0/0 | 0/0/0 | 0/0/0 | **1**/0/0 | 0/0/0 | 0/0/**1** | 0/0/**1** | 0/0/0 | **1**/0/**1** |
| Characidae | Jupiaba | *Jupiaba anteroides* | 0/0/0 | 0/0/0 | 0/0/0 | 0/0/0 | 0/0/0 | 0/0/0 | 0/0/0 | 0/0/0 | 0/0/0 | 0/0/0 | 0/0/**1** |
| Characidae | Knodus | *Knodus* sp. 1 | 0/0/0 | 0/0/0 | 0/0/0 | 0/0/0 | 0/0/0 | 0/0/0 | 0/0/0 | 0/0/0 | 0/0/0 | 0/0/0 | 0/0/**1** |
| Characidae | Knodus | *Knodus* sp. 2 | 0/0/0 | 0/0/0 | 0/0/0 | 0/0/0 | 0/0/0 | 0/0/0 | 0/0/0 | 0/0/0 | 0/0/0 | 0/0/0 | 0/0/**1** |
| Characidae | Knodus | *Knodus* sp. 3 | 0/0/0 | 0/0/0 | 0/0/0 | 0/0/0 | 0/0/0 | 0/0/0 | 0/0/0 | 0/0/0 | 0/0/0 | 0/0/0 | 0/0/**1** |
| Characidae | Knodus | *Knodus* sp. 4 | 0/0/0 | 0/0/0 | 0/0/0 | 0/0/0 | 0/0/0 | 0/0/0 | 0/0/0 | 0/0/0 | 0/0/0 | 0/0/0 | 0/0/**1** |
| Characidae | Knodus | *Knodus* sp. 5 | 0/0/0 | 0/0/**1** | 0/0/0 | 0/0/**1** | 0/0/0 | 0/0/0 | 0/0/0 | 0/0/0 | 0/0/0 | 0/0/**1** | 0/0/0 |
| Characidae | Knodus | *Knodus* sp. 6 | 0/0/0 | 0/0/0 | 0/0/0 | 0/0/0 | 0/0/**1** | 0/0/**1** | 0/0/**1** | 0/0/**1** | 0/0/0 | 0/0/0 | 0/0/0 |
| Characidae | Moenkhausia | *Moenkhausia bonita* | 0/0/0 | 0/0/0 | 0/0/0 | 0/0/0 | 0/0/0 | 0/0/0 | 0/0/0 | 0/0/0 | 0/0/0 | 0/0/0 | 0/0/**1** |
| Characidae | Moenkhausia | *Moenkhausia lepidura* | 0/0/0 | 0/0/0 | 0/0/0 | 0/0/**1** | 0/0/0 | 0/0/0 | 0/0/0 | 0/0/0 | 0/0/0 | 0/0/0 | 0/0/0 |
| Characidae | Moenkhausia | *Moenkhausia madeirae* | 0/0/0 | 0/0/0 | 0/0/0 | 0/0/0 | 0/0/**1** | 0/0/**1** | 0/0/**1** | 0/0/**1** | 0/0/**1** | 0/0/**1** | 0/0/**1** |
| Characidae | Moenkhausia | *Moenkhausia oligolepis* | 0/0/0 | 0/0/0 | 0/0/0 | 0/0/0 | 0/0/0 | 0/0/**1** | 0/0/0 | 0/0/0 | 0/0/0 | 0/0/0 | 0/0/0 |
| Characidae | Moenkhausia | *Moenkhausia sanctaefilomenae* | 0/0/0 | 0/0/0 | 0/0/0 | 0/0/0 | 0/0/0 | 0/0/0 | 0/0/0 | 0/0/0 | 0/0/0 | **1**/0/0 | **1**/0/0 |
| Characidae | Moenkhausia | *Moenkhausia* sp. | 0/0/0 | 0/0/0 | 0/0/0 | 0/**1**/0 | 0/0/0 | **1**/**1**/0 | 0/0/0 | **1**/**1**/0 | 0/**1**/0 | 0/**1**/0 | 0/**1**/0 |
| Characidae | Phenacogaster | *Phenacogaster* sp. | 0/0/0 | 0/0/0 | 0/0/0 | 0/0/0 | 0/0/0 | 0/0/0 | **1**/0/0 | **1**/0/0 | **1**/0/0 | **1**/0/0 | 0/0/0 |
| Characidae | Poptella | *Poptella compressa* | 0/0/0 | 0/0/0 | 0/0/0 | 0/0/0 | 0/0/0 | 0/0/0 | 0/0/0 | 0/0/0 | 0/0/0 | 0/0/0 | **1**/0/0 |
| Characidae | Roeboides | *Roeboides afinnis* | 0/0/0 | 0/0/0 | 0/0/0 | 0/0/0 | 0/0/0 | 0/0/0 | 0/0/**1** | 0/0/0 | 0/0/0 | 0/0/0 | 0/0/0 |
| Characidae | Roeboides | *Roeboides biserialis* | 0/0/0 | 0/0/0 | 0/0/0 | 0/0/0 | 0/0/0 | 0/0/0 | 0/0/0 | 0/0/**1** | 0/0/0 | 0/0/0 | 0/0/0 |
| Characidae | Roeboides | *Roeboides descalvadensis* | 0/0/0 | 0/0/0 | 0/0/0 | 0/0/0 | 0/0/0 | 0/0/0 | 0/**1**/0 | 0/**1**/0 | 0/**1**/0 | 0/0/0 | 0/0/0 |
| Characidae | Roeboides | *Roeboides* sp. | 0/0/0 | 0/0/0 | 0/0/0 | 0/0/0 | 0/0/0 | **1**/**1**/0 | 0/0/0 | **1**/**1**/0 | **1**/0/0 | **1**/0/0 | **1**/0/0 |
| Characidae | Serrapinnus | *Serrapinnus piaba* | **1**/0/0 | 0/0/0 | **1**/0/0 | **1**/0/0 | **1**/0/0 | **1**/0/0 | 0/0/0 | 0/0/0 | **1**/0/0 | 0/0/0 | **1**/0/0 |
| Characidae | Serrapinnus | *Serrapinnus* sp. 1 | 0/0/0 | 0/0/0 | 0/0/0 | 0/0/0 | 0/0/**1** | 0/0/**1** | 0/0/**1** | 0/0/**1** | 0/0/**1** | 0/0/**1** | 0/0/0 |
| Characidae | Serrapinnus | *Serrapinnus* sp. 2 | 0/0/0 | 0/0/0 | 0/0/0 | 0/0/0 | 0/0/0 | 0/0/0 | 0/0/**1** | 0/0/0 | 0/0/0 | 0/0/0 | 0/0/0 |
| Characidae | Tetragonopterus | *Tetragonopterus argenteus* | 0/0/0 | 0/0/0 | 0/0/0 | 0/0/0 | 0/0/0 | 0/0/0 | 0/0/0 | 0/**1**/**1** | 0/**1**/0 | 0/0/0 | 0/0/0 |
| Characidae | Tyttobrycon | *Tyttobrycon* sp. | 0/0/0 | 0/0/0 | 0/0/0 | 0/0/0 | 0/0/0 | 0/0/**1** | 0/0/0 | 0/0/0 | 0/0/0 | 0/0/0 | 0/0/0 |
| Cichlidae | Aequidens | *Aequidens tetramerus* | 0/0/0 | 0/0/**1** | 0/0/**1** | 0/0/0 | 0/0/0 | 0/0/0 | 0/0/0 | 0/0/0 | 0/0/0 | 0/0/0 | 0/0/0 |
| Cichlidae | Apistogramma | *Apistogramma* sp. | 0/0/0 | 0/0/0 | 0/0/0 | 0/0/0 | 0/0/0 | 0/0/0 | 0/0/0 | 0/0/**1** | 0/0/**1** | 0/0/0 | 0/0/**1** |
| Cichlidae | Bujurquina | *Bujurquina eurhinus* | 0/0/0 | 0/0/0 | 0/0/0 | 0/0/0 | 0/0/**1** | 0/0/**1** | 0/0/0 | 0/0/0 | 0/0/0 | 0/0/0 | 0/0/0 |
| Cichlidae | Cichlasoma | *Cichlasoma bimaculatum* | 0/**1**/0 | 0/0/0 | 0/**1**/0 | 0/**1**/0 | 0/**1**/0 | 0/**1**/0 | 0/**1**/0 | 0/**1**/0 | 0/**1**/0 | 0/**1**/0 | 0/**1**/0 |
| Cichlidae | Cichlasoma | *Cichlasoma boliviense* | 0/0/**1** | 0/0/0 | 0/0/0 | 0/0/**1** | 0/0/**1** | 0/0/0 | 0/0/**1** | 0/0/**1** | 0/0/0 | 0/0/0 | 0/0/0 |
| Cichlidae | Cichlasoma | *Cichlasoma* sp. | 0/0/0 | 0/0/0 | 0/0/0 | 0/**1**/0 | **1**/0/0 | **1**/0/0 | 0/0/0 | **1**/0/0 | **1**/0/0 | **1**/0/0 | **1**/0/0 |
| Cichlidae | Mesonauta | *Mesonauta festivus* | 0/0/0 | 0/0/0 | 0/0/0 | 0/0/0 | 0/0/0 | 0/0/0 | 0/0/0 | 0/0/0 | 0/**1**/0 | 0/0/**1** | 0/**1**/**1** |
| Cichlidae | Satanoperca | *Satanoperca jurupari* | 0/0/0 | 0/0/0 | 0/0/0 | 0/0/0 | 0/0/0 | 0/0/**1** | 0/0/0 | 0/0/**1** | 0/0/**1** | 0/0/**1** | 0/0/**1** |
| Cichlidae | Satanoperca | *Satanoperca* sp. | 0/0/0 | 0/0/0 | 0/0/0 | 0/0/0 | 0/0/0 | 0/**1**/0 | 0/0/0 | 0/**1**/0 | 0/**1**/0 | 0/**1**/0 | 0/**1**/0 |
| Cichlidae | Saxatilia | *Saxatilia lepidota* | **1**/0/0 | **1**/0/0 | **1**/0/0 | **1**/0/0 | **1**/0/0 | **1**/0/0 | 0/0/0 | **1**/0/0 | **1**/0/0 | **1**/0/0 | **1**/0/0 |
| Cichlidae | Saxatilia | *Saxatilia semicincta* | 0/0/**1** | 0/0/**1** | 0/0/**1** | 0/0/**1** | 0/0/0 | 0/0/**1** | 0/0/0 | 0/0/0 | 0/0/0 | 0/0/0 | 0/0/0 |
| Cichlidae | Saxatilia | *Saxatilia* sp. | 0/**1**/0 | 0/**1**/0 | 0/**1**/0 | 0/**1**/0 | 0/**1**/0 | 0/**1**/0 | 0/0/0 | 0/**1**/0 | 0/**1**/0 | 0/**1**/0 | 0/0/0 |
| Crenuchidae | Characidium | *Characidium* sp. | 0/0/0 | 0/0/0 | 0/0/0 | 0/0/0 | 0/0/0 | 0/0/0 | 0/0/**1** | 0/0/0 | 0/0/0 | 0/0/0 | 0/0/0 |
| Curimatidae | Curimatella | *Curimatella meyeri* | 0/0/0 | 0/0/0 | 0/0/0 | 0/0/0 | 0/0/0 | 0/0/0 | 0/0/0 | 0/0/**1** | 0/0/0 | 0/0/0 | 0/0/0 |
| Curimatidae | Curimatella | *Curimatella* sp. | 0/0/0 | 0/0/0 | 0/0/0 | 0/0/0 | 0/0/0 | 0/0/0 | 0/**1**/0 | 0/0/0 | 0/**1**/0 | 0/0/0 | 0/0/0 |
| Curimatidae | Cyphocharax | *Cyphocharax* sp. | 0/0/0 | 0/0/0 | 0/0/0 | 0/0/0 | 0/0/0 | **1**/**1**/0 | **1**/0/0 | **1**/**1**/0 | **1**/**1**/0 | **1**/**1**/0 | **1**/**1**/0 |
| Curimatidae | Cyphocharax | *Cyphocharax spiluropsis* | 0/0/0 | 0/0/0 | 0/0/0 | 0/0/0 | 0/0/0 | 0/0/**1** | 0/0/0 | 0/0/**1** | 0/0/0 | 0/0/**1** | 0/0/0 |
| Curimatidae | Potamorhina | *Potamorhina altamazonica* | 0/0/0 | 0/**1**/0 | 0/0/0 | 0/0/0 | 0/**1**/0 | 0/**1**/**1** | 0/**1**/**1** | 0/**1**/**1** | 0/**1**/**1** | 0/**1**/**1** | 0/**1**/0 |
| Curimatidae | Psectrogaster | *Psectrogaster rutiloides* | 0/0/0 | 0/0/0 | 0/0/0 | 0/0/0 | 0/0/**1** | 0/0/0 | 0/**1**/0 | 0/0/**1** | 0/0/0 | 0/**1**/0 | 0/0/0 |
| Curimatidae | Steindachnerina | *Steindachnerina* aff. *dobula* | 0/0/0 | 0/0/0 | 0/0/0 | 0/0/0 | 0/0/**1** | 0/0/**1** | 0/0/0 | 0/0/**1** | 0/0/**1** | 0/0/0 | 0/0/0 |
| Curimatidae | Steindachnerina | *Steindachnerina guentheri* | 0/0/0 | 0/0/0 | 0/0/0 | 0/0/0 | 0/0/**1** | 0/**1**/**1** | 0/0/0 | 0/0/0 | 0/0/0 | 0/**1**/**1** | 0/0/0 |
| Curimatidae | Steindachnerina | *Steindachnerina* sp. | 0/0/0 | 0/0/0 | 0/0/0 | 0/0/0 | 0/0/0 | 0/0/0 | 0/0/0 | 0/0/**1** | 0/0/0 | 0/0/0 | 0/0/0 |
| Cynodontidae | Cynodon | *Cynodon gibbus* | 0/0/0 | 0/0/0 | 0/0/0 | 0/0/0 | 0/0/0 | 0/0/0 | 0/0/0 | 0/0/**1** | 0/0/0 | 0/0/0 | 0/0/0 |
| Cynodontidae | Cynodon | *Cynodon* sp. | 0/0/0 | 0/0/0 | 0/0/0 | 0/0/0 | 0/0/0 | 0/0/0 | 0/0/0 | 0/0/0 | 0/0/0 | 0/**1**/0 | 0/0/0 |
| Cynodontidae | Hydrolycus | *Hydrolycus scomberoides* | 0/0/0 | 0/0/0 | 0/0/0 | 0/0/0 | 0/0/0 | 0/0/0 | 0/0/0 | **1**/0/0 | 0/0/0 | 0/0/0 | 0/0/0 |
| Erythrinidae | Hoplerythrinus | *Hoplerythrinus unitaeniatus* | 0/0/**1** | 0/0/0 | 0/**1**/**1** | 0/0/0 | 0/0/0 | 0/**1**/0 | 0/0/0 | 0/0/0 | 0/0/0 | 0/0/0 | 0/0/0 |
| Erythrinidae | Hoplias | *Hoplias malabaricus* | **1**/**1**/0 | 0/0/**1** | 0/0/**1** | **1**/0/**1** | **1**/**1**/**1** | **1**/**1**/**1** | **1**/**1**/**1** | **1**/**1**/**1** | **1**/**1**/**1** | **1**/**1**/**1** | **1**/**1**/0 |
| Gymnotidae | Gymnotus | *Gymnotus carapo* | 0/0/0 | 0/0/0 | 0/0/0 | 0/0/0 | 0/0/0 | **1**/**1**/0 | **1**/**1**/0 | **1**/0/0 | **1**/**1**/0 | **1**/**1**/0 | **1**/0/0 |
| Hemiodontidae | Anodus | *Anodus elongatus* | 0/0/0 | 0/0/0 | 0/0/0 | 0/0/0 | 0/0/0 | 0/0/0 | 0/0/0 | 0/0/**1** | 0/0/0 | 0/0/0 | 0/0/0 |
| Heptapteridae | Pimelodella | *Pimelodella* sp. | 0/0/0 | 0/0/0 | 0/0/0 | 0/0/0 | 0/0/0 | 0/0/0 | 0/0/0 | **1**/0/0 | 0/0/0 | 0/0/0 | 0/0/0 |
| Hypopomidae | Brachyhypopomus | *Brachyhypopomus pinnicaudatus* | 0/0/0 | 0/0/0 | 0/0/0 | 0/0/0 | 0/0/0 | 0/**1**/0 | 0/0/0 | 0/0/0 | 0/0/0 | 0/0/0 | 0/0/0 |
| Hypopomidae | Brachyhypopomus | *Brachyhypopomus* sp. | 0/0/0 | 0/0/0 | 0/0/0 | 0/0/0 | 0/0/0 | **1**/0/0 | 0/0/0 | 0/0/0 | 0/0/0 | **1**/0/0 | **1**/0/0 |
| Iguanodectidae | Bryconops | *Bryconops* sp. | 0/0/0 | 0/0/0 | **1**/0/0 | 0/0/0 | 0/0/0 | 0/0/0 | 0/0/0 | 0/0/0 | 0/0/0 | 0/0/0 | 0/0/0 |
| Iguanodectidae | Bryconops | *Bryconops melanurus* | 0/0/0 | 0/0/0 | 0/0/0 | 0/0/**1** | 0/0/0 | 0/0/0 | 0/0/0 | 0/0/0 | 0/0/0 | 0/0/0 | 0/0/0 |
| Loricariidae | Ancistrus | *Ancistrus sp.* | 0/0/0 | 0/0/0 | 0/0/0 | 0/0/0 | 0/0/0 | **1**/**1**/0 | 0/0/0 | **1**/0/0 | **1**/**1**/0 | **1**/**1**/0 | **1**/**1**/0 |
| Loricariidae | Hypoptopoma | *Hypoptopoma* cf. *bianale* | 0/0/0 | 0/0/0 | 0/0/0 | 0/0/0 | 0/0/0 | 0/0/0 | 0/0/0 | 0/0/0 | 0/0/**1** | 0/0/0 | 0/0/0 |
| Loricariidae | Hypoptopoma | *Hypoptopoma gulare* | 0/0/0 | 0/0/0 | 0/0/0 | 0/0/0 | 0/0/0 | 0/**1**/0 | 0/**1**/0 | 0/**1**/0 | 0/0/0 | 0/0/0 | 0/**1**/**1** |
| Loricariidae | Hypoptopoma | *Hypoptopoma* sp. | 0/0/0 | 0/0/0 | 0/0/0 | 0/0/0 | 0/0/0 | **1**/0/0 | 0/0/0 | **1**/0/0 | **1**/0/0 | **1**/0/0 | **1**/0/0 |
| Loricariidae | Hypostomus | *Hypostomus* sp. | 0/0/0 | **1**/0/0 | 0/0/0 | 0/0/0 | **1**/**1**/0 | **1**/**1**/0 | **1**/**1**/0 | **1**/**1**/0 | **1**/**1**/0 | **1**/**1**/0 | **1**/**1**/0 |
| Loricariidae | Hypostomus | *Hypostomus* sp.1 | 0/0/0 | 0/0/0 | 0/0/0 | 0/0/0 | 0/0/0 | 0/0/**1** | 0/0/**1** | 0/0/**1** | 0/0/**1** | 0/0/**1** | 0/0/0 |
| Loricariidae | Hypostomus | *Hypostomus* sp.2 | 0/0/0 | 0/0/0 | 0/0/0 | 0/0/0 | 0/0/0 | 0/0/0 | 0/0/**1** | 0/0/**1** | 0/0/0 | 0/0/0 | 0/0/0 |
| Loricariidae | Loricariichthys | *Loricariichthys platymetopon* | 0/0/0 | 0/0/0 | 0/0/0 | 0/0/0 | **1**/**1**/**1** | **1**/**1**/**1** | **1**/**1**/0 | **1**/**1**/**1** | **1**/**1**/0 | **1**/**1**/0 | **1**/**1**/**1** |
| Loricariidae | Loricariichthys | *Loricariichthys* sp. | 0/0/0 | 0/0/0 | 0/0/0 | 0/0/0 | 0/0/0 | 0/0/0 | 0/0/0 | 0/0/0 | 0/0/0 | 0/0/**1** | 0/0/0 |
| Loricariidae | Pterygoplichthys | *Pterygoplichthys disjunctivus* | 0/0/0 | 0/0/0 | 0/0/0 | 0/0/0 | 0/0/0 | 0/0/0 | 0/0/**1** | 0/0/0 | 0/0/0 | 0/0/0 | 0/0/0 |
| Pimelodidae | Hemisorubim | *Hemisorubim platyrhynchos* | 0/0/0 | 0/0/0 | 0/0/0 | 0/0/0 | 0/0/0 | 0/0/0 | 0/0/0 | **1**/0/0 | 0/0/0 | 0/0/0 | 0/0/0 |
| Pimelodidae | Hypophthalmus | *Hypophthalmus edentatus* | 0/0/0 | 0/0/0 | 0/0/0 | 0/0/0 | 0/0/0 | **1**/0/0 | 0/0/0 | 0/0/0 | 0/0/0 | 0/0/0 | **1**/0/**1** |
| Pimelodidae | Hypophthalmus | *Hypophthalmus* sp. | 0/0/0 | 0/0/0 | 0/0/0 | 0/0/0 | 0/0/0 | 0/**1**/0 | 0/0/0 | 0/0/0 | 0/0/0 | 0/0/0 | 0/**1**/0 |
| Pimelodidae | Pimelodus | *Pimelodus* sp. | 0/0/0 | 0/0/0 | 0/0/0 | 0/0/0 | 0/0/0 | 0/0/0 | **1**/0/0 | **1**/0/0 | **1**/0/0 | **1**/0/0 | 0/0/0 |
| Pimelodidae | Pimelodus | *Pimelodus tetramerus* | 0/0/0 | 0/0/0 | 0/0/0 | 0/0/0 | 0/0/0 | 0/0/0 | 0/0/0 | 0/**1**/0 | 0/**1**/0 | 0/0/0 | 0/0/0 |
| Pimelodidae | Pseudoplatystoma | *Pseudoplatystoma* sp. | 0/0/0 | 0/0/0 | 0/0/0 | 0/0/0 | 0/0/0 | 0/0/0 | **1**/0/0 | 0/0/0 | 0/0/0 | 0/0/0 | 0/0/0 |
| Pimelodidae | Pseudoplatystoma | *Pseudoplatystoma tigrinum* | 0/0/0 | 0/0/0 | 0/0/0 | 0/0/0 | 0/0/0 | 0/0/0 | **1**/0/0 | **1**/0/0 | **1**/0/0 | 0/0/0 | 0/0/0 |
| Pimelodidae | Sorubim | *Sorubim lima* | 0/0/0 | 0/0/0 | 0/0/0 | 0/0/0 | 0/0/0 | 0/0/0 | **1**/**1**/0 | **1**/0/**1** | 0/0/0 | 0/0/0 | 0/0/0 |
| Prochilodontidae | Prochilodus | *Prochilodus lineatus* | 0/0/0 | **1**/0/0 | 0/0/0 | **1**/0/0 | **1**/0/0 | 0/0/0 | **1**/0/0 | **1**/0/0 | **1**/0/0 | **1**/0/0 | **1**/0/0 |
| Prochilodontidae | Prochilodus | *Prochilodus nigricans* | 0/0/0 | 0/0/0 | 0/0/0 | 0/0/0 | 0/0/0 | 0/0/**1** | 0/**1**/**1** | 0/**1**/**1** | 0/**1**/0 | 0/**1**/**1** | 0/**1**/0 |
| Serrasalmidae | Mylossoma | *Mylossoma albiscopum* | 0/0/0 | 0/0/0 | 0/0/0 | 0/0/0 | 0/0/0 | 0/0/0 | 0/0/0 | 0/**1**/**1** | 0/0/0 | 0/0/0 | 0/0/0 |
| Serrasalmidae | Serrasalmus | *Serrasalmus maculatus* | 0/0/0 | 0/0/0 | 0/0/0 | 0/0/0 | 0/0/0 | 0/0/**1** | 0/0/**1** | 0/0/**1** | 0/0/0 | 0/0/**1** | 0/0/**1** |
| Serrasalmidae | Serrasalmus | *Serrasalmus rhombeus* | 0/0/0 | 0/0/0 | 0/0/0 | 0/0/0 | 0/0/0 | 0/**1**/**1** | 0/0/**1** | 0/0/0 | 0/0/0 | 0/**1**/0 | 0/**1**/**1** |
| Serrasalmidae | Serrasalmus | *Serrasalmus* sp. | 0/0/0 | 0/0/0 | 0/0/0 | 0/0/0 | 0/0/0 | **1**/0/0 | **1**/0/0 | **1**/0/0 | **1**/0/0 | **1**/0/0 | **1**/0/0 |
| Serrasalmidae | Serrasalmus | *Serrasalmus spilopleura* | 0/0/0 | 0/0/0 | 0/0/0 | 0/0/0 | 0/0/0 | 0/0/0 | 0/0/0 | 0/0/**1** | 0/0/0 | 0/0/0 | 0/0/**1** |
| Sternopygidae | Eigenmannia | *Eigenmannia* cf. *macrops* | 0/0/0 | 0/0/0 | 0/0/0 | 0/0/0 | 0/0/0 | 0/0/0 | 0/0/**1** | 0/0/0 | 0/0/0 | 0/0/0 | 0/0/0 |
| Sternopygidae | Eigenmannia | *Eigenmannia* gr. *trilineata* | 0/0/0 | 0/0/0 | 0/0/0 | 0/0/0 | 0/0/0 | 0/0/0 | 0/0/0 | 0/0/0 | 0/0/0 | 0/**1**/0 | 0/0/0 |
| Sternopygidae | Eigenmannia | *Eigenmannia limbata* | 0/0/0 | 0/0/0 | 0/0/0 | 0/0/0 | 0/0/0 | 0/0/0 | 0/0/0 | 0/0/0 | 0/0/0 | 0/**1**/0 | 0/0/0 |
| Sternopygidae | Eigenmannia | *Eigenmannia* sp. | 0/0/0 | 0/0/0 | 0/0/0 | 0/0/0 | 0/0/0 | **1**/**1**/0 | **1**/**1**/0 | 0/0/**1** | **1**/0/0 | 0/0/0 | 0/0/0 |
| Sternopygidae | Eigenmannia | *Eigenmannia virescens* | 0/0/0 | 0/0/0 | 0/0/0 | 0/0/0 | 0/0/0 | 0/0/**1** | 0/0/0 | 0/0/0 | 0/0/0 | 0/0/0 | 0/0/0 |
| Sternopygidae | Sternopygus | *Sternopygus macrurus* | 0/0/0 | 0/0/0 | **1**/**1**/0 | **1**/**1**/**1** | 0/0/0 | **1**/0/0 | **1**/**1**/0 | **1**/**1**/**1** | **1**/**1**/0 | **1**/**1**/0 | **1**/0/0 |
| Synbranchidae | Synbranchus | *Synbranchus marmoratus* | 0/0/0 | 0/0/0 | 0/0/0 | 0/0/0 | 0/0/0 | 0/0/0 | 0/0/0 | 0/0/0 | 0/0/0 | **1**/0/**1** | 0/0/0 |
| Triportheidae | Triportheus | *Triportheus angulatus* | 0/0/0 | 0/0/0 | 0/0/0 | 0/0/0 | 0/0/0 | 0/0/0 | 0/0/**1** | 0/0/**1** | 0/0/0 | 0/0/**1** | 0/0/**1** |
| Triportheidae | Triportheus | *Triportheus rotundatus* | 0/0/0 | 0/0/0 | 0/0/0 | 0/0/0 | 0/0/**1** | 0/0/0 | 0/0/0 | 0/0/0 | 0/0/0 | 0/0/0 | 0/0/0 |
| Triportheidae | Triportheus | *Triportheus* sp. | 0/0/0 | 0/0/0 | 0/0/0 | 0/0/0 | 0/0/0 | 0/**1**/0 | 0/**1**/0 | 0/**1**/0 | 0/**1**/0 | 0/**1**/0 | 0/**1**/0 |
